# Open-source robotic chip-to-plate interface for high-throughput microfluidic generation of materials libraries

**DOI:** 10.64898/2026.05.12.724546

**Authors:** Isabel B. Navarro, Gregory Datto, Lameck Beni, Diego Barragan, Katherine J Mossburg, Shiny Shen, Andrew R. Hanna, David P. Cormode, David Issadore

## Abstract

Data-driven materials development requires large, well-characterized libraries of precisely defined formulations. While microfluidic platforms excel at generating highly controlled materials, their throughput is often limited by the challenge of efficiently interfacing device outputs with standard well plates. This bottleneck frequently necessitates manual transfer or non-microfluidic workflows, constraining both throughput and reproducibility. Here, we present LMNOP-bot (Libraries of Micro- and Nano-materials, OPen-source bot), an open-source robotic platform for the automated generation and collection of micro- and nanomaterial libraries from serial microfluidic outputs. Using synchronized, pressure-driven flow, LMNOP-bot enables continuous formulation and direct deposition into standard well plates. The system is low-cost (<$700, excluding pressure regulators), constructed from readily available or easily fabricated components, and designed for broad accessibility. LMNOP-bot collects ≥30 µL per formulation at a rate of one sample every four seconds, representing an approximately 50× increase in throughput over existing serial microfluidic workflows, and operates robustly for over 10,000 runs without maintenance. We demonstrate compatibility with both PDMS/glass and commercial polycarbonate devices, with seamless interfacing to 96- and 384-well plates. Repeated sampling confirms high precision and reproducibility. By removing a key bottleneck in microfluidic library generation, LMNOP-bot enables rapid, scalable, and accessible exploration of material design spaces.

## Introduction

The precise control of flows at the micrometer scale achievable in microfluidic chips has made possible the generation of micro- and nano-materials with highly-controlled, precisely-defined structures and functions suitable for medicine and biological research that are not attainable with conventional bulk techniques^1-3^. However, structure–function relationships are often unknown for the micro-/nano-scale materials generated by microfluidics, leaving formulation and material selection to be guided largely by empirical exploration. Identifying materials and formulations for particular applications often requires searching a high-dimensional parameter space, encompassing constituent components, their proportions, and their physical combination within a device, motivating researchers to generate and screen libraries of many formulations^4-6^, especially as machine-learning–driven and autonomous discovery approaches become more popular^7-11^. There has been significant recent progress in designing microfluidic systems specifically for library generation in materials research, leveraging small dead volumes, rapid response in flow rates, and automation, which results in reduced equipment and staffing needs, and improved throughput compared to bulk methods^1, 12-16^. For complex micro- and nano-materials, where there may be simultaneous development of new components and of formulations using those components, as well as subsequent characterization of the formulated final products, it’s often the microfluidic generation step which hinders high-throughput material discovery^4, 6, 7, 17, 18^, with researchers turning to robotic or manual pipetting for higher-throughput nanoparticle formulation, even where microfluidic generation is superior^6^. While very-large-scale microfluidic integration (VLSMI) can define ≥10^4^ generators on a single chip, originally developed to increase throughput of a single material by operating identical units in parallel^2^, a major limitation arises when attempting to scale up the number of distinct formulations, such as in a library. Application of parallel, non-identical generators have recently been demonstrated to significantly increase the rate of unique formulation production^19^. To be compatible with downstream analyses, a material library generator must either collect formulations separately, or label them so they can be separated and identified. While barcoding can be applied to some use cases^14, 20^, not all materials or downstream assays are compatible with existing labelling schemes; for example, barcoding by fluorescence uses spectra that might be needed for assay readout. Thus, each generator requires a corresponding outlet, creating a bottleneck. The challenge is no longer fitting more generators on a chip, but rather efficiently collecting and managing the output from numerous outlets.

While great progress has been made in using microfluidics to generate libraries of materials, current methods of microfluidic material library generation are primarily serial, from a single device and outlet, rather than parallelized, due to this difficult tradeoff between formulation and volume throughputs^21, 23^. In situ microarrays can be defined by single-nozzle printing, requiring on the order of 10 hours per 1500 combinations^13, 24-27^. While the position in the array holds identity information, just as in a well-plate, such 2-D libraries are only compatible with *in situ* assays, which characterize the materials on the surface where they were deposited, limiting their applicability^13^. Methods for separating single-outlet libraries of liquid-phase products include serial collection, either manually or by commercial robotics^11, 21, 28^. Non-microfluidic generation of libraries of self-assembling nanoparticles via pipetting robots, as well as in situ microarray printing, have both demonstrated the potential of robotics to increase the efficiency of micro-/nano-material library generation^5, 23, 29^. Recently, we demonstrated robotically automated lipid nanoparticle (LNP) library generation and collection from a multi-outlet parallelized microfluidic chip^19^, our approach to addressing microfluidic library formulation scalability. The synchronized robotic and microfluidic system was custom-built for high-throughput collection of LNPs into a 96-well plate from the 2×4 outlet array of a silicon/glass chip. This system was able to generate LNPs at a rate of 1 formulation per 4s, ∼100× faster than conventional microfluidic approaches, and about 50× faster than some commercial LNP library systems^19, 22^. This previous work was designed solely for silicon/glass devices; in this work, we apply the concept to more common PDMS/glass and polycarbonate devices.

In this paper, building on our prior work, we present an automated, generalizable, modular robotic chip-to-world system for library generation and collection, which synchronizes a well-plate compatible X/Y/Z motion platform with automated pressure-driven fluidics in order to produce plates of microfluidically-generated materials of varying formulations (Figure 1). All hardware and electronic components are standard and readily available from suppliers such as McMaster-Carr and Digikey, or included as design files for printed circuit board (PCB) manufacturing, laser cutting, or 3D printing on the LMNOP-bot GitHub‡. The platform is designed for modular integration across a broad range of microfluidic architectures to create an accessible entry point to high-throughput library generation for research laboratories. We refer to this system as LMNOP-bot (Libraries of microfluidically-generated Micro- and Nanomaterials, OPen-source bot). Despite its relatively low-cost construction (<$700), hobbyist-grade electronics, and 3D-printed parts, we achieve sufficient precision to allow for collection into 96- and 384-well plates, withstanding 10,000 cycles of wear and tear; compatibility with standard well plate formats is a desirable feature for high-throughput library generation for compatibility with downstream assays and other laboratory automation equipment^6, 18, 23, 30^. Our previous work employed a multi-generator, multi-outlet silicon/glass chip; here, we demonstrate modular compatibility of the robotic interface with a single-generator PDMS/glass chip, which is a more common, simple, and inexpensive fabrication method, as well as with a commercially available polycarbonate chip. To demonstrate the precision of our system to define ratios of fluid inputs, we measured replicates of 192 distinct combinations of three dyes with water dilution. Finally, to provide an example use case, we generated and measured silver sulfide nanoparticles, a promising X-ray imaging contrast agent^31, 32^, and found them to be comparable to gold standard displacement-driven microfluidically-generated, bulk-collected particles. By demonstrating robust mechanical and fluidic accuracy and precision, with 50× more formulations per hour than commercial automation systems^22^, and broad flexibility in the device used, we demonstrate the value of this platform to the research community.

**Figure 1.**
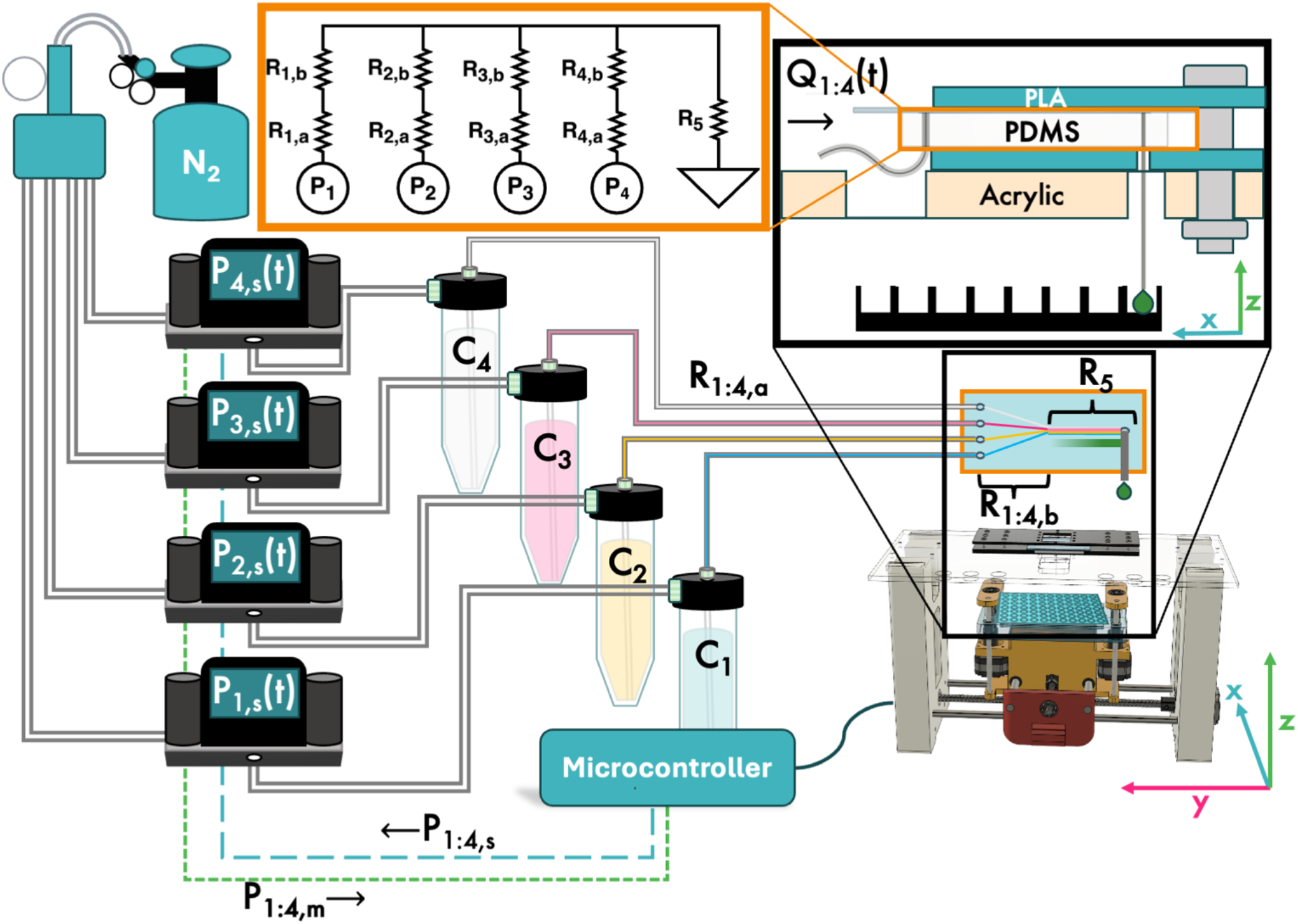
Overview of LMNOP-bot. Nitrogen tank supplies pressure to four pressure regulators, which recieve the set pressure, P_s_(t), from a microcontroller, and return the measured pressure, P_m_(t). Pressures P_1:4, m_ enter component reservoirs C_1:4_. Tubing leaving the reservoirs contributes resistances R_1:4,a_, while microfluidic channels in a PDMS chip contribute further resistances R_1:4,b_, and the exit channel and outlet needle contribute shared resistance R_5_. The chip is sandwiched between two layers of a simple chip holder which screws into an acrylic top plate. The microcontroller actuates motion in X, Y, and Z directions in order to collect product from the chip into a well plate.

## Materials and Methods

### Mechanical design of robotic system

The design of LMNOP-bot centers around the X/Y/Z manipulation of a standard format well plate, synchronized with the precise manipulation of pressures (Fig. 1). A 3D-printed tray into which these plates fit snugly is at its core. The majority of parts were made from fused filament fabrication in polyethylene terephthalate glycol (PETG) or carbon fiber-blended polylactic acid (PLA-CF) filaments, printed on a Bambu Lab P1S, with 0.4mm brass nozzle. Load-bearing custom lead screw nuts were fabricated from Durable resin on a Form 2 SLA printer (Formlabs, USA). The motion of this tray in all directions is performed by closed-loop NEMA-17 stepper motors (17HS15-1504-ME1K, StepperOnline, Canada) driving 1″/rotation lead screws (99030A100, McMaster-Carr, USA) to move platforms along linear rails, with two synchronized motors lifting the tray in the Z direction. Each platform integrates a lead screw nut, which can be either 3D-printed or purchased (95075A101, McMaster-Carr), to translate rotational motion of the motor to linear motion of the plate. Detailed assembly and use instructions, including a complete bill of materials, design files, and sample code, are contained in the LMNOP-bot Github repository. A laser-cut acrylic sheet provides a surface to affix microfluidic chip holders above the collection plate; this was made by shaping a ¼″ nominal thickness acrylic sheet with a laser cutter (PLS6.150D, Universal Laser Systems, USA). The large surface area allows for addition of various useful through-holes, which can be used for mounting a tube microscope or, as we have, holding 15mL Falcon tubes (Fig. 1).

### Pressure-driven fluidics

For fluidic performance experiments, pressure-driven flow was modulated with the use of up to four pressure regulators (PCD-30PSIG-D-PCV30/5P, Alicat, USA), connected to 2-port aluminum caps (LVF-KPT-S-2, Darwin Microfluidics, France) on 15-mL Falcon tubes. All tubing in and out of these caps had a 1/16″ OD, held in place with ¼-28 threaded flangeless nuts and ferrules (P-230 and P-200, IDEX Health & Science, USA). The pressure regulators were fed from a 5-port manifold (P-155, IDEX Health & Science) connected to the nitrogen cylinder, with inlet pressure modulated to ≅30 PSI (slightly above the maximum pressure to be used for any single component) by two analog pressure regulating valves. A tolerance threshold of ±0.3PSI was used to indicate when the pressure had equilibriated after changing the set pressure.

### Synchronization

LMNOP-bot is controlled by an Arduino microcontroller (Arduino MEGA 2560 Rev3, Arduino, Italy), which synchronizes changes in the set pressures, P_s_, measured pressures, P_m_, and plate location (Fig. 2) through the pressure regulators (PCD-30PSIG-D-PCV30/5P, RIN, 5IN, RANGE (50 PSIG) P1: 100 PSIG, P2: ATM, VOL: 15 mL, Alicat, USA) and motor drivers (A4988 Stepper Motor Driver Carrier, #1182, Pololu, USA). An example of the sequence of motions and synchronized pressure changes is shown in Figure 3: first, the plate is lifted such that the microfluidic device outlet(s) enter a purge well(s); concurrently, a set pressure (or list of set pressures) is sent to the pressure regulators, which begin shifting towards those pressures. After the time necessary for the measured pressure(s) to come within a tolerance threshold of the set pressure(s), plus a stabilization time, the plate is lowered away from the needles and moved so that the outlet is aligned with the next well/set of wells. At this point, P_m_ has stabilized to ≅P_s_, and the plate is raised so that the outlets are within the collection well(s). After a predetermined collection period, the plate is lowered such that the outlet needles are clear of the plate, the plate is moved to the next purge well, and the cycle starts again.

**Figure 2.**
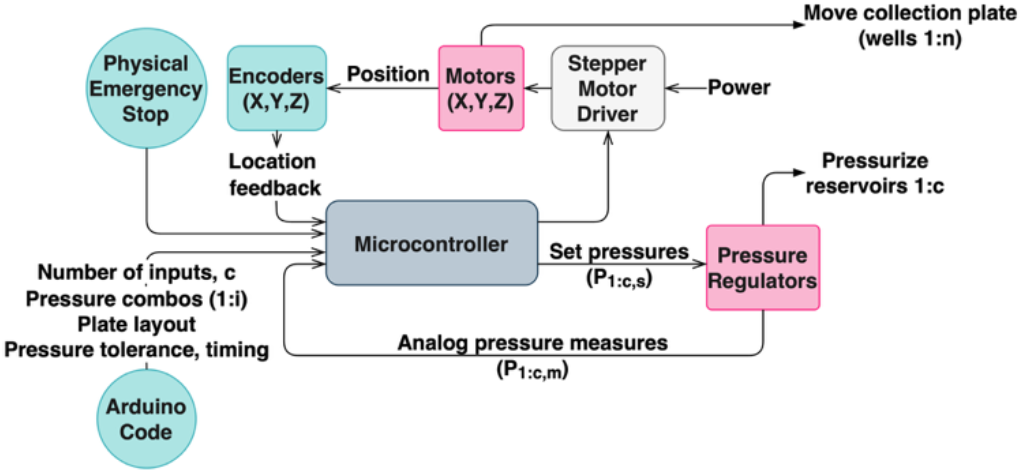
Control of synchronized motion and pressure. Arduino code is used to set the collection plate layout in terms of rows, columns, and spacing, the number of component inputs, the timing and tolerance preferences, and the desired pressure combinations. Once started, the Arduino controls both stepper motors and pressure regulators in order to generate and collect the desired formulations.

**Figure 3.**
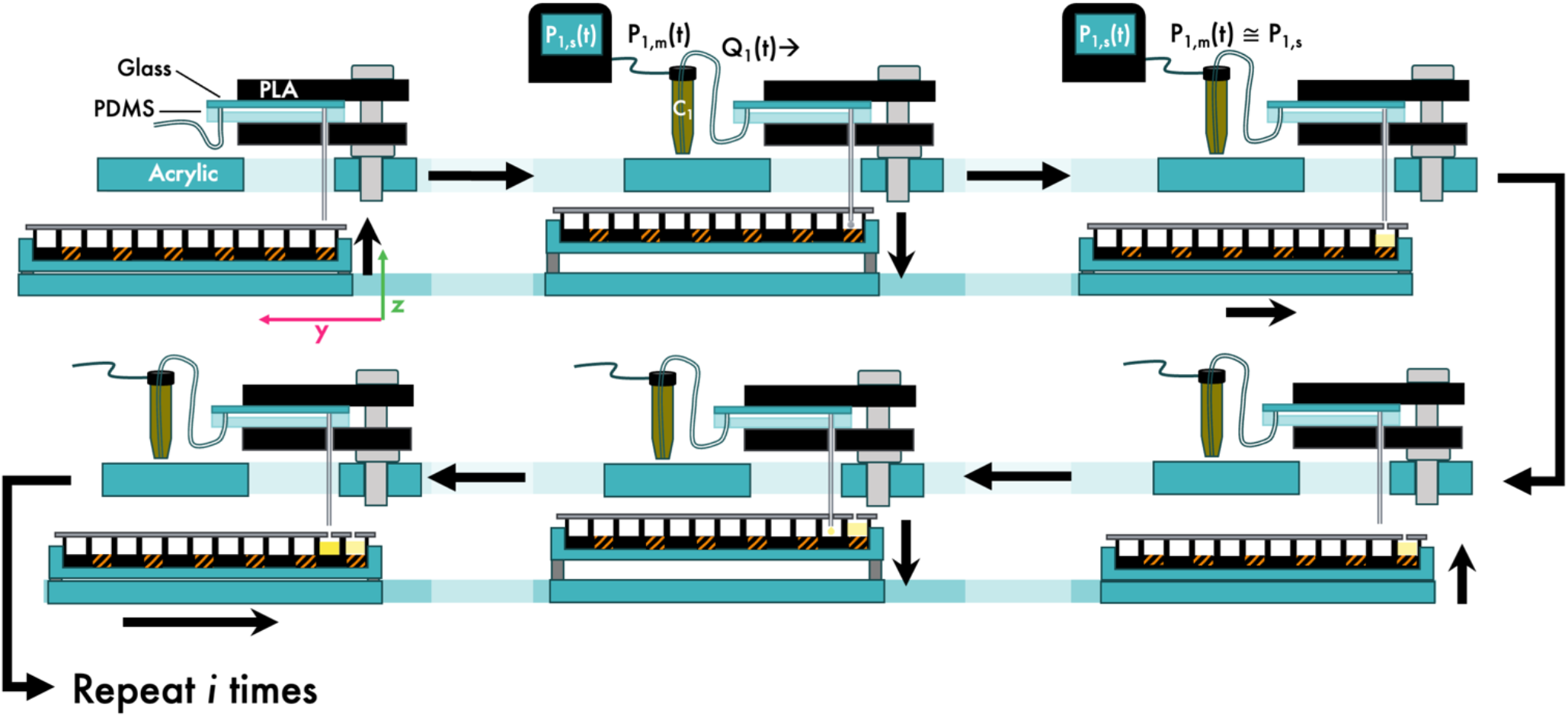
Synchronization of plate motion with pressure regulation. Once the device outlet is within the purge well (diagonal stripes), a set pressure (P_s_) is sent to the regulator. The plate is not retracted until the measured pressure P_m_ is ≅ P_s_ to within a set tolerance. The plate is then moved such that the device outlet enters a collection well to collect the product of the desired pressure settings. Once the set collection time has elapsed, the plate is again retracted and will move to the next purge well, at which point a new pressure setting will be sent.

The path taken between collection wells depends on the arrangement of the outlet(s) and on the type of plate used. A path which visits every well in a horizontal serpentine is the default, and can be easily modified to suit experimental needs by modifying the number of rows and columns, and the well spacing (ESI Code Supplement). In this paper, different paths are used for each experiment: visitation of a 2×4 outlet array to 96- and 384-well plates for motion precision experiments, and visitation of a single outlet to 384- and 96-well plates for dye collection and silver sulfide nanoparticle generation, respectively (ESI Fig. 1).

### Interface for benchmarking location precision

For motion precision testing, we employed a custom aluminum chip holder designed for a 2×4 array of outlets (ESI Fig. 2). Each threaded outlet was interfaced with a Luer adapter (P-619, IDEX, USA), which held in place a 9mm long 30G needle (LDS-30009I, TSK Laboratory America, USA). In order to measure the accuracy of repeated movements, a foil film (AB0626, ThermoFisher, USA) was applied over the plate being used. Each puncture of the needle was thus ‘recorded’ as a hole in the foil, such that overlapping punctures appear as a single hole, which was carefully removed from the plate and affixed to a labelled index card for ease of handling and storage. Sub-millimeter-scale holes were imaged microscopically, with multi-frame images stitched using FIJI^33^, while macroscopic holes were imaged by a phone camera equipped with a macro lens, using a sub-millimeter ruler (TOL-15295, SparkFun Electronics, USA) for scale.

### Interface for PDMS chip fluidic characterization

For characterization of fluidic performance, we used a 4-inlet, 1-outlet PDMS/glass chip with 200×95µm rectangular channels, for an equivalent hydraulic diameter of ≅129µm. Each inlet was formed by a 1.5mm biopsy punch, and directly interfaced to the Falcon tube reservoirs with 14” of .007” inner diameter (ID) PTFE tubing (1/16” outer diameter (OD), 49210-07-C, MicroSolv, USA), creating a resistor for each inlet upstream of the PDMS chip (Figure 1). The outlet of the chip was a 1.5” 21G needle, which was separated from its hub by soaking overnight in acetone. This was fitted into an outlet hole punched in the PDMS with a .75mm biopsy punch. This assembly was held in place over the plate-handling stage by a simple 3D-printed sandwich (Fig. 1) held in place by screws which fit into alignment holes in the acrylic top plate of the bot. The port and needle are centered over a hole in the 3D-printed base plate. Unlike many custom device holders which require a certain relative position of outlets and inlets, this alignment system’s latitude and multiple clamping locations allows for the use of multi-device chip formats, as different devices could be accessed simply by removing the outlet needle, shifting the chip, and reinserting the outlet needle and inlet tubing. Products were collected into and had absorbance measured in a flat- and clear-bottom, 384-well plate (781096, Greiner Bio-One, USA).

### Interface for silver sulfide nanoparticle generation

Our method for the generation of Ag_2_S (silver sulfide) nanoparticles is similar to previous reporting^34^. As the flow of silver nitrate was to be varied by an order of magnitude, while the flow of sodium sulfide was to be varied by a factor of <2, two different types of tubing were chosen to give different resistances: 16” of 300µm ID, 1/16” OD polytetrafluoroethylene (PTFE) tubing (49225-80, Microsolv, USA) and 14” of .007” ID tubing, respectively. A simple platform was designed and 3D printed to hold a commercially available polycarbonate herringbone mixer device (Fluidic 187, Microfluidic ChipShop, Germany) by friction fit over the well plate (ESI Fig. 3). The tubing was interfaced with the chip using Tube Tuck connectors for 1/16” tubing, and the same needle used for dye experiments was interfaced as an outlet using a Tube Tuck for 1/32” tubing (Fluidic 1581 and 1580, Microfluidic ChipShop, Germany).

### Benchmarking location precision

Initial investigations on the precision and robustness of motion control measured the relative area of punctures from 10 laps (ESI Fig. 1A) of a 96-well plate employing 0, 500, and 1000ms of optional delay (Fig. 4) between X/Y motion and subsequent puncture by motion in the Z-axis; the areas of the 10 overlapping punctures were recorded on a separate foil for each delay time (Fig. 5A). LMNOP-bot was then run for 10,000 laps without optional delay, with the punctures again recorded into a foil (Fig. 5D). Finally, 10 laps’ worth of punctures were recorded for each of the delay time settings to determine whether the 10,000 runs had reduced the precision (Fig 5A). Our testing of the effect of 10,000 laps of wear and tear on the bot made use of 3D-printed lead screw nuts made with Durable resin (Durable Resin V2, Formlabs, USA) on a Form2 printer.

**Figure 4.**
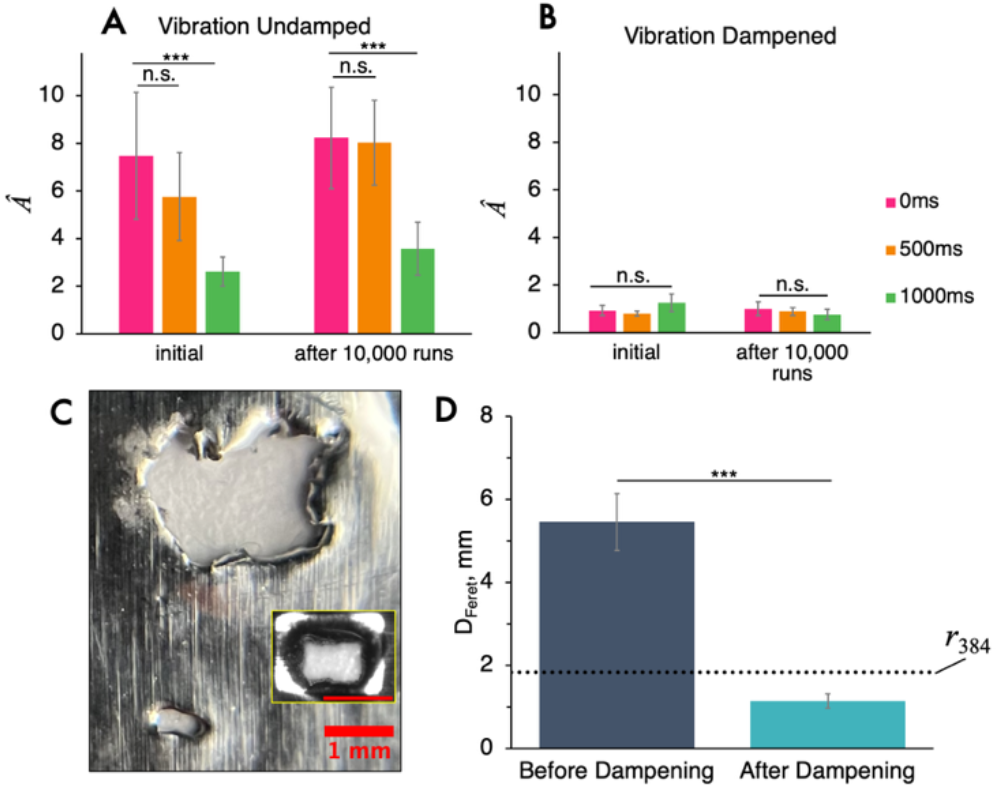
*Â* is hole area divided by the cross-sectional area of 300µm needle. A. Comparison of the area of 10 sequential punctures on a 96-well plate, at three levels of optional delay, before and after running 10,000 laps, normalized by the cross-sectional area of 300µm OD needle; n=10. B. Comparison of the area of 10 sequential punctures on a 384-well plate, before and after running 10,000 laps, normalized by the cross-sectional area of the needle; n=40. C. Representative image of 10,000-puncture cluster before dampening vibrations, taken with phone camera using macro lens. Inset: microscope image of 10,000-puncture cluster after addressing sources of vibration. Scale bars 1mm. D. Feret diameter of 10,000-puncture cluster created with no optional delay with and without vibration-dampening measures. Dotted line indicates typical inner diameter of 384-well plate wells (1.825mm).

**Figure 5.**
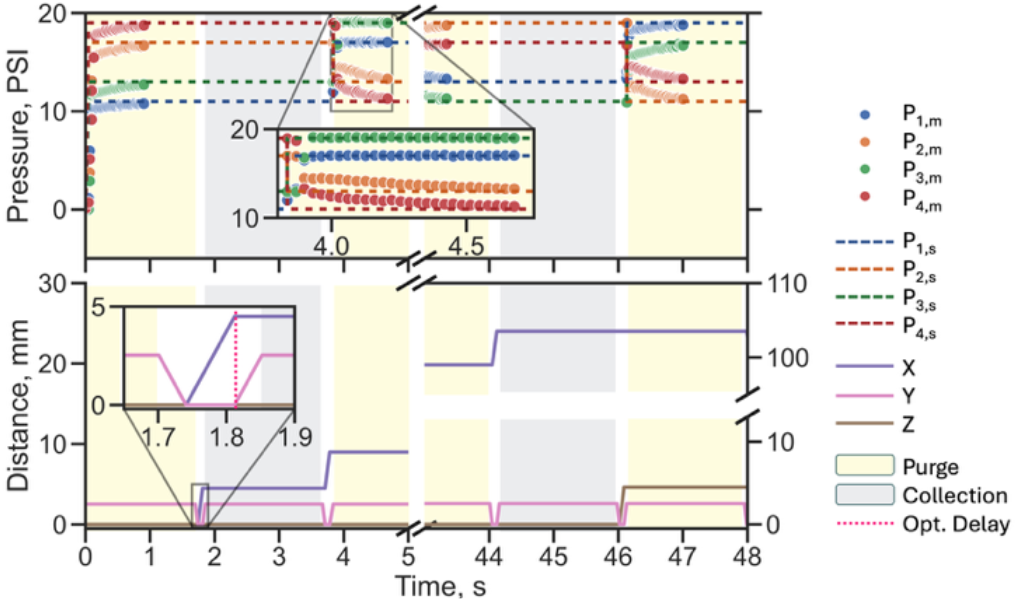
Synchronization of motion and pressure control, showing optional delay timing, pressure stabilization, and purge vs. collection timing. Pressure was measured once per 32ms.

### Fluidic characterization via mixing of three dyes

Batch dilutions of three dyes (Patent Blue, Red Dye 40, and McCormick® Yellow Food Color^§^ were loaded into the reservoirs as inputs (components, c) to the chip, with the fourth input being water. Calibration curves (ESI Fig. 4) were created in parallel with dilution in order to choose optimized dye dilutions for which the expected volumes of 1–16µL would span most of the available absorbance range. An absorbance calibration curve was made from each dye dilution at 637, 490, and 425nm (for Patent Blue, Red Dye 40, and Yellow dye, respectively) in order to later extrapolate the composition of collected mixtures. 192 different pressure combinations were chosen to create 192 unique mixtures of dye and water, with pressures selected from the prime numbers between 11 and 23 PSI, inclusive. The formulations were collected for 1.6s each into a clear-bottom 384-well polystyrene microplate (781096, Greiner Bio-One, USA). Because the volume collected was not the same for each pressure combination, a daughter plate was prepared by transferring 30µL of each collection well by multichannel pipette, in order to determine the proportion of the volume contributed from the water input. Absorbances were measured on a plate reader (Tecan Infinite 200 Pro M Plex, Switzerland). Signals from calibration dilutions were fit to a two parameter linear function. The average signal of the triplicate blank was subtracted from all points at each wavelength. Estimates of dye dilution volumes were calculated from their signals using the calibration curves after subtracting the average blank signal. The same pressure settings were run 3 times, refilling the 15mL reservoirs from the batch dilutions, and inserting a new collection plate between runs.

### Silver sulfide nanoparticle generation

Silver sulfide nanoparticles were generated from the microfluidic mixing of an aqueous solution made with 75mL of water with 42.5mg of silver nitrate and 2.5mmol of 2-mercaptopropionic acid added, titrated to pH 7.5 with sodium hydroxide, and an aqueous solution prepared by adding 5mg of sodium sulfide to 25mL of water at room temperature (all reagents from Sigma-Aldrich, USA). Due to the volume needed for downstream transmission electron microscopy (TEM) analysis, each of 8 formulations was collected into 11 wells of a 96-well plate, with the first well of each row used for equilibration. In order to stabilize the particles^35^, each collection well was pre-filled with 175µL of HPLC-grade water. Thus the total pooled volume for each replicate was ≅3mL. Once the particles were collected, the collection from the pressure setting corresponding to the 3:1 flow ratio was pooled for each plate, and concentrated using 3kDa MW cutoff filtration tubes (MAP003C38, Pall Corporation, USA) in preparation for TEM imaging. Pressures were selected to center around a 3:1 silver nitrate solution:sodium sulfide solution flow rate ratio, with a 0.5mL/min flow of sodium sulfide. A new custom chip holder was designed and 3D printed from PETG, held in place with aligned screw-holes cut into a new piece of top acrylic (ESI Fig. 3).

## Results and Discussion

### Design

The aim was to develop a custom plate-handling robot that simultaneously controlled motion and pressure regulation. We modelled this robot against commercially-available computer numerical control (CNC) and 3D printing machines. The torque of stepper motors is reduced at higher speeds, so we selected a maximum speed of ≅200rpm based on the stepper motors’ speed-torque curve to maintain optimal torque. Due to this limitation, we opted for a 1 ″/rotation lead screw, which would allow for translation across 2 wells of a 96-well plate in <0.2s, minimizing potential contamination (supported by ESI Supplemental Video). The laser-cut ¼” acrylic top plate can be customized to include cutouts for the reservoirs, guides for the tubing, and mounting holes for custom chip holders as well as portable digital microscopes, which would be useful for autonomous optimization^11^. So long as the first outlet is aligned with a preset location, it is trivial to design a new device holder and acrylic top plate combination (ESI Fig. 2). The acrylic top plate rests on 3D-printed uprights with a close but non-joining fit, allowing for displacement, rather than compression, in the case that device outlets hit the bottom of the well, or are misaligned, protecting fragile microfluidic devices during testing. A new holder for the commercial chip used for sulfide nanoparticle generation was devised and fabricated in just a few hours using 3D printing. The automating platform of the bot can be assembled at a cost of <$700, with compatible pressure regulators available for ≅$1500/input. While we chose to develop a custom PCB for portability and reliability, many iterations of LMNOP-bot used simple perma-proto boards (590, Adafruit, USA) to connect the electronics; however, the PCB design is included in the design files (ESI Fig. 5). The available travel distance is 11.5cm in the Y direction, 12cm in the X direction, and importantly, 3.5cm in the Z direction, which allows for compatibility with both standard and deep-well plates. To encourage and facilitate replication of this system, the supplemental GitHub repository^‡^ includes the relevant computer-assisted design (CAD) files, PCB design, complete bills of materials, detailed assembly and operation instructions, and required code for control of the platform.

The LMNOP-bot’s motion and pressure systems are coordinated by our open-source C++ code for the Arduino MEGA (GitHub and ESI Code Supplement). The code is adjustable for various plate formats and can dynamically scale for up to nine inputs. Users can specify the number of rows and columns, the spacing, and the number of input regulators to control, as well as the stabilization and collection times to suit their application.

### Location precision

Initially, we saw a significant improvement in precision (Fig. 4) with the addition of a one-second delay between lateral and vertical motion; one second, but not 500ms, was sufficient time for the vibration to significantly diminish. However, this level of motion delay would add over 1.5 minutes to the collection of 48 unique formulations from a single generator, and >6 minutes to collection from a single generator into a 384-well plate. When we ran the bot for 10,000 laps with no delay, the Feret diameter of the 10,000-hole cluster was over 5mm, indicating that the precision without delay would not be acceptable for repeated use with 384-well plates, which have a 4.5mm center-to-center spacing (Fig 4C, D). Based on the results of this initial testing, it was determined that undamped vibrations were causing a level of imprecision that would preclude the use of 384-well plates. To optimize precision and speed, we minimized the impact of vibration found in the prior experiments in three ways: by increasing the length of the linear bearings; by reducing the tolerance of the 3D-printed lead screw nuts; and by using LocTite Threadlocker Blue 242 (209728, Henkel Corp., USA) on the coupling set screws to prevent catastrophic slipping, which could lead to a one-time offset within the run (Figure 5C, ESI Fig. 6). Increasing the length of the lead screw nuts nominally increases their price. After these improvements, we repeated the collection of a 10-lap baseline, 10,000-run durability test, and the post-test 10-lap cluster. The normalized area, *Â*, (equal to the area of the hold divided by the cross-sectional area of the needle) of the post-test cluster, without optional delay, was ≤1— that is, the area of the hole was smaller than the needle making them. In these cases, the punctures totally overlapped, leaving a D-, rather than O-shaped hole, due to the bevel of the needle (Figure 4B; ESI Fig 7B). Additionally, the average Feret diameter of the 10,000-hole cluster after minimizing the impact of vibration was 1.1mm, indicating that the improved LMNOP-bot would be suitable for frequent use with 384-well plates (the wells of which have a radius of 1.55mm) (Fig. 4D). Finally, the mechanical precision of the bot was comparable before and after 10,000 runs, indicating that the mechanical apparatus, including the 3D-printed lead screw nuts, is robust enough for frequent operation. Use of four 3D-printed lead screw nuts constitutes a savings of nearly $140 in parts, while maintaining the necessary accuracy and performance after 10,000 runs.

**Figure 6.**
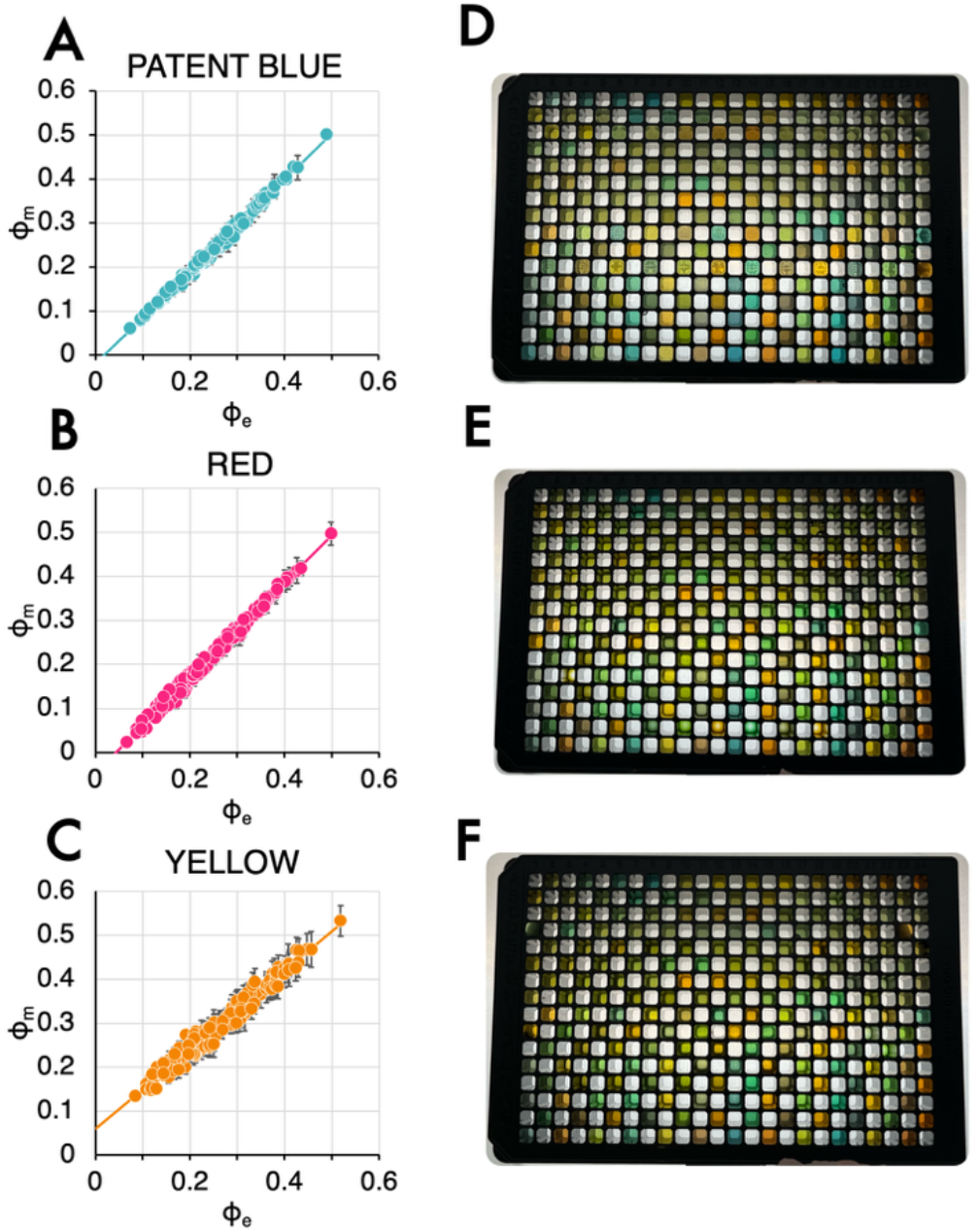
Measured vs. expected flow proportion (ϕ_m_ vs. ϕ_e_) of Patent Blue, Allura Red, and Yellow dye. The slopes of the best-fit lines are 1.042, 1.083, and 0.899; the intercepts are -0.019, -0.047, and 0.059, respectively. The R^2^ values are 0.996, 0.993, and 0.968, respectively. Error bars are 95% CI.D-F are underlit images of replicates 1-3 of 30µL aliquots of the 192 distinct formulations. Absorbances were read immediately after generation and aliquot transfer by plate reader; these images were taken after the water had evaporated during storage and the plates were subsequently reconstituted and thus are for illustrative purposes only. Well spacing is a standard 4.5mm center-to-center.

**Figure 7.**
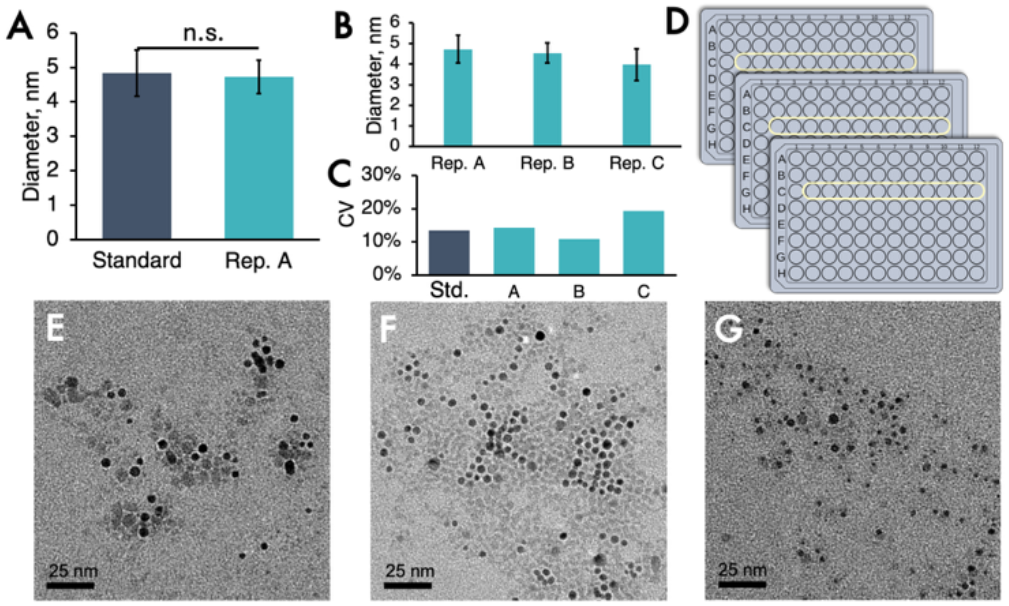
A: Average diameter of 100 Ag_2_S nanoparticles measured by TEM, from the gold-standard displacement-driven bulk collection compared to pressure-driven robotic collection, Replicate A. Error bars 95% CI. B: Average diameter of 100 nanoparticles measured by TEM, from LMNOP-bot–generated replicates A–C. Error bars are one standard deviation. C: Coefficient of variation for nanoparticle diameter for displacement-driven bulk collection and pressure-driven robotic collection replicates A–C. D: Collection across 11 wells pooled for each replicate. E–G: Representative TEM micrographs from replicates A-C.

### Fluidic performance

The tubing ID and length were chosen such that the resistance created by the tubing, *R*_*a*_, (Fig. 1) would be approximately an order of magnitude higher than the on-chip resistance *R*_*b*_, and could achieve the desired flow rates using pressures accessible with our 50PSI max pressure regulators, while maintaining a pressure exiting the tubing of ≤10PSI, in order to minimize the deformation of the PDMS device. While tubing provides an accessible method for increasing resistance, lithographically-defined on-chip resistors can provide an increased precision in flow rate^19^. This resistance allowed for relatively linear response to pressure changes despite low on-chip resistance between inlets, making this interface suitable for a wide range of chip geometries, including those not initially designed for pressure-driven flow (ESI Fig. 8).

Collection of ≥30µL of dye mixtures, using a pressure tolerance of ±0.3PSI, an 800ms stabilization time, and 1600ms collection time led to a formulation rate of ≥1 formulation per 4 seconds when collecting into the 384-well plate (Fig. 5). Three plates of 192 formulations were produced in <45 minutes, inclusive of refilling reservoirs, transferring collected plates to a plate reader, and placing a new plate into LMNOP-bot, a rate comparable to the readout of the three dye absorbances via plate reader. While automated or integrated plate switching and transfer may be desirable for future self-driving lab applications8, it does not appear to save an appreciable amount of time for use even with an automated downstream characterization that requires no additional reagents or processing.

Supplying a higher pressure than necessary to the regulators may increase the time needed for equilibriation; thus we recommend supplying slightly more than the maximum pressure setting that will be requested of any one regulator. Equilibriation within the 0.3PSI threshold was consistently reached in <1s (Fig. 5). Use of a static gun such as the Milty Zerostat 3 (Goldring, England) was necessary to dissipate electrostatic charge on the polycarbonate well-plates used for collection; otherwise, small aqueous droplets respond to the electrostatic force and can enter wells other than the one the outlet is positioned over (ESI Video).

The expected flow rates of each component (Q_c,e_) were calculated for each set of pressures by solving the resistance network shown in Figure 1. ϕ_*c,e*_ (Fig. 6), the estimated proportion of flow for each component, *c*, was calculated as

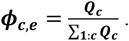

The measured flow proportion, ϕ_*c,m*_ was determined for each pressure setting from absorbance measurements on a 30µL aliquot collected at that pressure setting (Fig. 6, D-F), combined with prepared calibration curves **(SI Fig. 4)**, to give

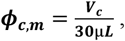

where *V*_*c*_ is the volume of component *c*.

Predicted values agreed well with observed values across all three dyes (root mean squared error = 1.14%, 2.93%, and 3.67% for Patent Blue, Allura Red, and Yellow dye, respectively across triplicate measurements), with R^2^ values ≥0.97 for all components (Fig. 6).

A few main sources of interexperimental variation were identified in the fit of measured flow rate ratios vs. expected flow rate ratios. First, there was some variation in replicate points for the calibration curves (SI Fig. 4), which may be due to imprecision in preparation of the calibration curves. This variance compounds when calculating the amount of Yellow dye, as both Allura Red and and FD&C Yellow 5 are present in the Yellow dye mixture. An additional and potentially significant source of error is the application of a pressure tolerance; specifically, we chose a 0.3PSI tolerance in conjunction with an 800ms stabilization time. In Figure 5, we can see two behaviors for Pm: a near-immediate shift to ∼P_s_ (inset, P_1,m_ and P_3,m_), or a significant but insufficient shift, followed by gradual approach to P_s_ (inset, P_2,m_ and P_4,m_). While we expect that the pressure would continue to approach the true P_s_ during the stabilization time, we did not continue collect Pm measurements during this time. If the P_m_ sometimes remained at the outer limits of the pressure tolerance, and othertimes approached P_s_, this would contribute ≤3% pressure difference between runs. Additionally, =examination of the individual residuals (ESI Fig. 9) show a reduction in residual magnitude in the second run only discernable in the measured Yellow flow. The first run was performed on one day; the second and third runs were performed on the following day. This transient experimental artifact suggests there are factors yet to be optimized, such as refreshing reservoirs between runs, which could further improve the performance and predictability of the LMNOP-bot system.

This imperfect fit is also evident in the slopes (≠1) and intercepts (≠0) of our best fit lines. This was due in part to our use of a simplified geometric model to estimate the resistances of our microfluidic chip. Alternatives, such as COMSOL modelling, or more simply, the use of this dye measurement experiment to calibrate our resistance model, could strengthen the correlation. Nevertheless, prediction of results from our simple model was strongly linear, precise, and calibratable, showing the value of this method.

### Silver sulfide nanoparticle collection and characterization

Silver sulfide (Ag_2_S) nanoparticles are an area of active medical imaging research owing to their promise as contrast agents for X-ray imaging in dense breast tissue_31, 36_. As size is known to impact relevant properties, such as organ partitioning and excretion, microfluidics are appealing due to their low heterogeneity, but still-limited by the low number of formulations generated per hour^32^. To demonstrate the flexibility and utility of LMNOP-bot for material library generation, Ag_2_S nanoparticles were made with a commercially available polycarbonate device and measured by TEM using a Talos L120C Transmission Electron Microscope as previously described^32^ (Fig. 7, E-G). As above, the necessary tubing resistance was determined from the range of flow rates needed, the pressure supply possible using our regulators, and a maximum outlet pressure from the tubing near 10PSI. We compare the particle size of our first collection to a typical displacement-driven generation, of which 100mL is typically collected. 11 wells of a 96-well plate were pooled in order to supply sufficient material for processing; due to the serial generation requiring less start-up volume, even this required 100× less volume than manual syringe-pump methods for the 8 formulations made. Using TEM, the measured diameters of our three automated collections were 4.7±.6nm, 4.5±.7nm, and 4.0±.8nm, compared to 4.8±.7nm for the gold standard displacement-driven microfluidic generation with bulk collection. The size and standard deviation observed are comparable to expected literature values^32^.

## Discussion

Looking to future integration with existing open-source microfluidic hardware and lab automation, we expect that the cost of pressure regulators could be significantly reduced by adoption of open-source Arduino-controlled pressure regulators^37^. This would be valuable to researchers seeking to control a large number of inputs, as the cost of a single regulator in our presented system is double the fixed cost of the base automation system. The addition of automated input switching^38^ would greatly increase the value of this bot for library generation. A recent multi-input parallelized library generation scheme was reported which draws from a well-plate loaded with FACS machine-readable barcodes, loading each parallel generator with unique dyes which label the formulation; the barcodes can later be used to separate the single outlet stream of products via FACS into an on-plate library^14^. However, this method limits the spectra available for characterization assays, and necessitates the FACS sorting step before any assay can be run. Furthermore, all inputs in this system are subject to the same input pressures, leading to the same formulations with different components, with each new pressure combination requiring a reset of the pressure chamber and manual relocation of the outlets into new collection wells. Implementing even a single switchable component input with LMNOP-bot could open a much-needed new dimension of high-throughput automated library generation. Switching inputs may necessitate addition of a washing step to remove traces of the earlier compound, but this waste could be collected in the current purge wells. Another useful extension would be the attachment of a simple microscope to observe the microfluidic chip, which could give video feedback to adjust material generation parameters^11, 39^.

One significant source of time savings not captured by the time per formulation is the time required for reagent preparation or pre-mixing. In the Sunscreen (Unchained Labs) commercial plate-to-plate LNP formulation platform22, two plates with pre-mixed componenets (internal and external) are the input. Without a pipetting robot, one can imagine their preparation adds significant additional manual labor, time, and sources of error. The serial generation of products from continuous flow inputs in LMNOP-bot removes this time and labor-intensive process.

In this work, we controlled pressure to control flow rates, collecting for a constant amount of time, leading to variable collected volumes. Implementing variable collection times could simplify downstream assays by allowing for collection of consistent volumes regardless of flow rate. Estimating the flow rate from geometrically-derived resistances can limit the accuracy of flow rate predictions for pressure-driven flow through compliant chips. Inline flow meters, *post hoc* volume measurements^14^, or dye measurement experiments such as the one undertaken here, can be used to empirically improve resistance estimations. There is limited space for variables on the microcontroller; thus, there may be a trade-off between the precision of collection times and the precision of the pressure settings, especially for protocols with higher numbers of distinct formulations.

We did not observe fouling of the PDMS/glass or polycarbonate microfluidic devices. However, this was likely due in large part to the fact that we mixed only small molecule food dyes and aqueous dilutions, which are highly water soluble. Other components may interact with the walls and with each other in such a way as to aggregate and cause fouling and clogging; the severity may also be impacted by device geometry^40-42^. Washing steps^43^, surface coatings^42^, cleaning techniques^44^, and changes to device geometry^40^ are all supported methods to reverse, reduce, and prevent fouling. We recommend that users continue to use any anti-fouling techniques necessary for normal device function; one advantage of the LMNOP-bot system is the reduced volumes necessary for start-up may allow for similar volumetric throughput between a plate of library production and single-formulation batch collection, leading to comparable fouling over comparable volumetric output.

## Conclusions

We offer an automated microfluidic library generation and collection platform, which is capable of collecting ≥30µL of one formulation per four seconds from a single-generator device, a 50× improvement over commercially available microfluidic automation platforms^22^, and is flexibly compatible with both custom PDMS/glass and commercial polycarbonate devices. Compared to such plate-to-plate platforms, the serial generation of formulations directly from components also saves the time needed for pre-formulation mixing of the desired component ratios. The platform is widely adaptable and could be interfaced with robotic arms or cameras for low-cost self-driving laboratory setups. We show that it is compatible with 96- and 384-well plates, and robust enough, even with 3D-printed parts, for over 10,000 uses without maintenance. The automation platform can be assembled at a cost of <$700, exclusive of pressure regulators. This technology could be extended by an input-switching mechanism to automate generation of complex libraries of formulations. By reducing the labor required for high-throughput library generation, we envision that LMNOP-bot will enable microfluidic material researchers to more easily generate, collect, and characterize libraries of new materials, and aid in developing the datasets necessary for machine-learning–guided material discovery.

## Supporting information

Supplemental Arduino Code

## Author contributions

I.B.N. and D.A.I. conceptualized the robotic collection system. D.P.C., D.B., and I.B.N. conceptualized silver sulfide experiments. I.B.N. led mechanical design, conceptualized and performed validation experiments, and analyzed and interpreted data. I.B.N., G.D., and L.B. developed the system hardware and electronics; I.B.N. and G.D. developed the control code. A.H. designed the 2×4 outlet array and microfluidic device used for dye mixing. S.S. developed assembly methods and documentation and was responsible for bot assembly and initial validation. D.B. guided silver sulfide nanoparticle experiments, and K.J.M. performed associated experiments; D.B. performed associated data analysis. I.B.N. prepared and edited the manuscript. All authors provided feedback on the manuscript. D.A.I. and D.P.C. provided funding, resources, mentorship, and project supervision.

## Data availability

All code, design files, bills of material, and instructions for reproducing and using the LMNOP-bot are hosted at the project GitHub found at www.github.com/IssadoreLab/LMNOP-bot. Additionally, the control code is included in the ESI of this article.

## Acknowledgements

The authors gratefully acknowledge use of facilities and instrumentation supported by NSF through the University of Pennsylvania Materials Research Science and Engineering Center (MRSEC) (DMR-2309043) as well as Cooperative Agreement DBI – 2400135. We also acknowledge funding support from the NIH (R21-CA297703, R01-CA291880).

## Notes

^§^The ingredients of McCormick® Yellow Food Dye are: Water, propylene Glycol, FD&C Yellow 5, 0.1% Propylparaben, *and FD&C Red 40* (i.e, Allura Red). This will subsequently be referred to as Yellow; calibration curves (SI Fig. 4) were used to account for the increased absorbance in the Allura Red channel.

