## Supplementary material for "Open-source robotic chip-to-plate interface for high-throughput microfluidic generation of materials libraries": SI Figures

### Supplemental Information and Figures for Open-source robotic chip-to-plate interface for high-throughput microfluidic generation of materials libraries

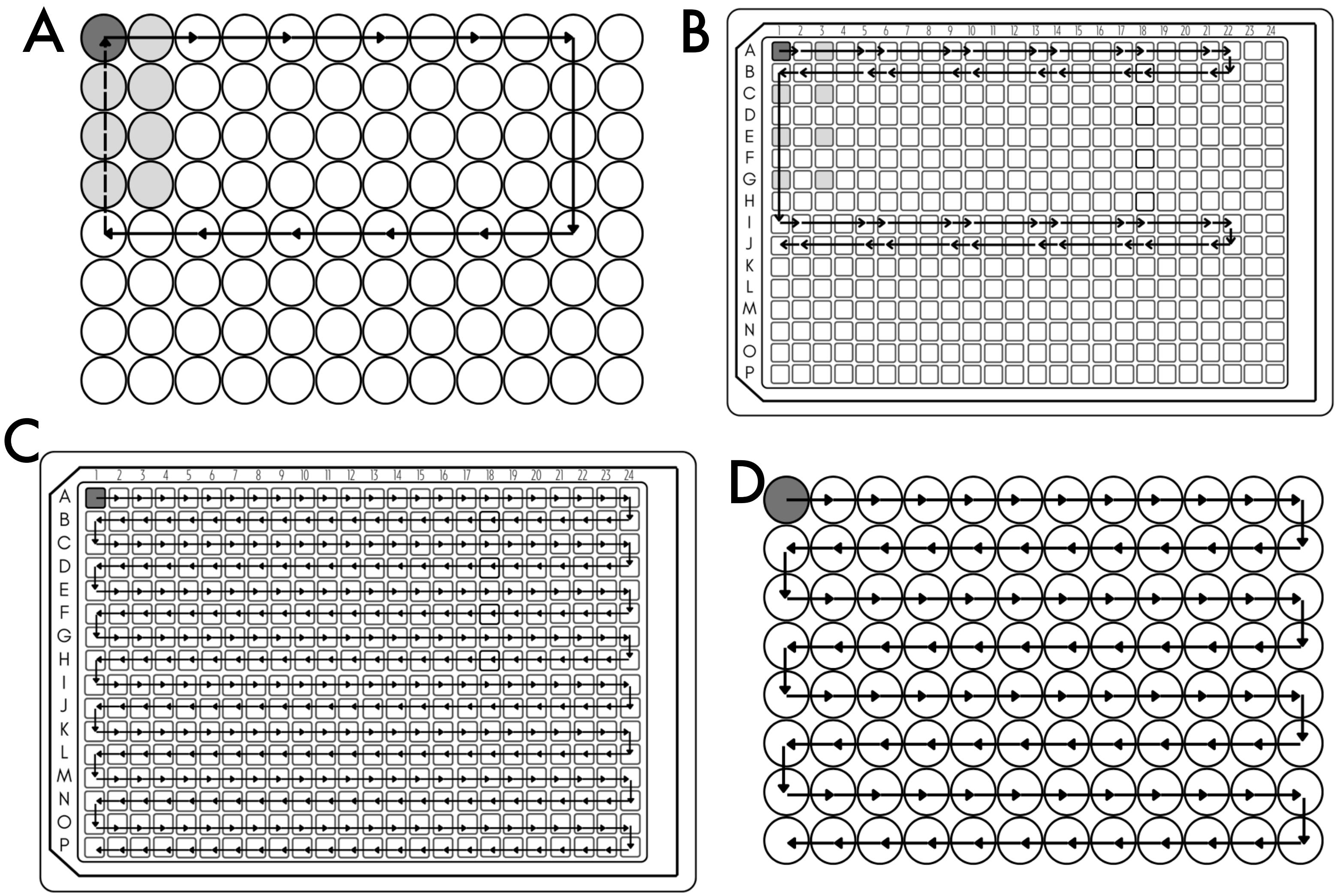


**SI Figure 1.** Paths used to traverse the collection plate. All paths start with the upper left "A1" needle in well A1; arrow heads indicate the well "A1" needle stops in if array of outlets is used—otherwise, indicates where single outlet stops.
A: 2x4 outlet array, 96-well plate
B: 2x4 outlet array, 384-well plate
C: single outlet, 384-well plate
D: single outlet, 96-well plate.

Starting location of "A1" needle or single outlet needle

Starting location of remaining 2x4 array outlet needles

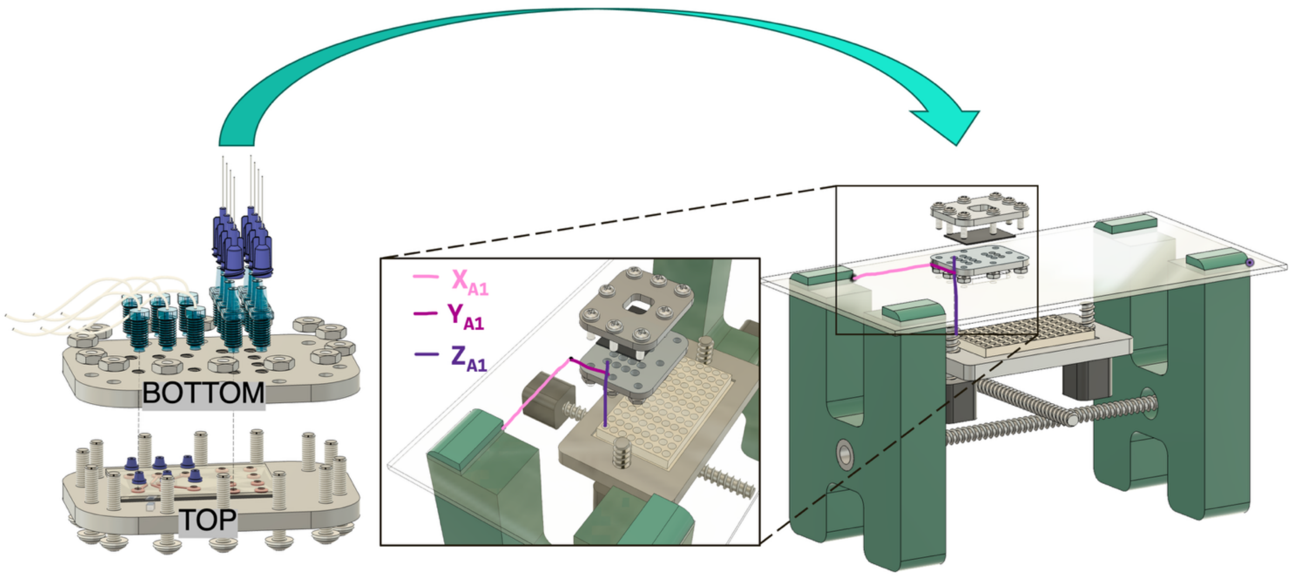


**SI Figure 2**: Adaptation of the acrylic plate and custom 3D-printed chip holder to other layouts is simple. So long as outputs are aligned to a standard well-plate array size (such as 9mm center-to-center spacing, used for 96-well layouts, shown here), one simply ensures that the top-right outlet aligns with the preset location of the first well, which can be adjusted in the code by the variables `center_x` and `center_y`. By maintaining these relative distances XA1, YA1, and ZA1, as annotated in the file for the laser-cut acrylic top, it is trivial to design a new device holder for your application.

**SI Figure 3: Chip holder for commercial polycarbonate chip.**

This chip holder/interface was designed around the commercially available polycarbonate chip (Fluidic 187, Microfluidic ChipShop) by taking measurements with a caliper. On the second design iteration, the chip was held satisfactorily by press fit around the protruding inlets and outlets.

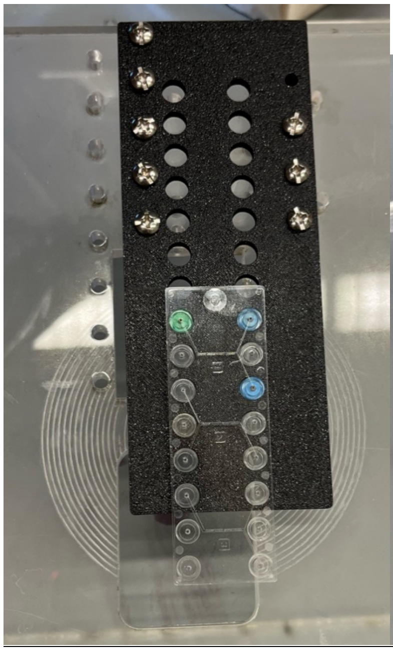

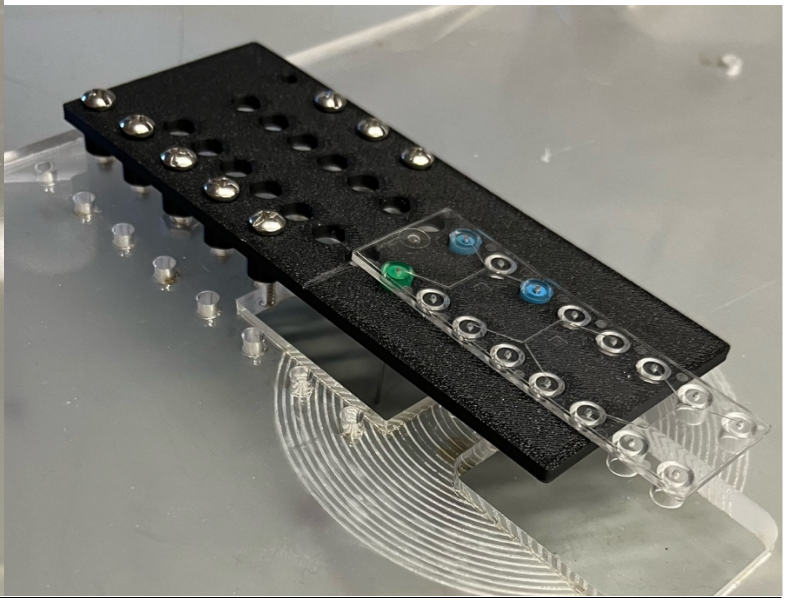

View from above View from above side

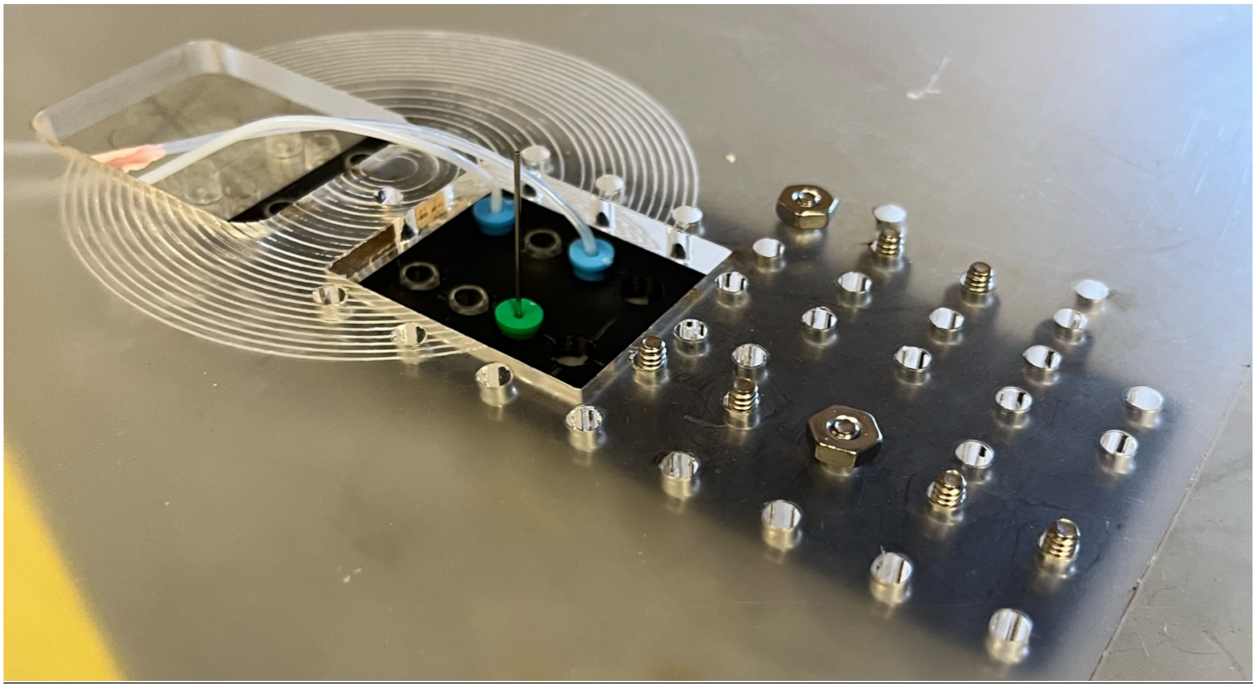

View from below side showing tube tuck Luer connectors

**SI Figure 4:** Calibration curves. Dilutions were prepared in conjunction with trialling calibration curves such that the expected volumes (1-16μL) spanned most of the available absorbance range (0-4). Error bars are 95% CI from n=3.

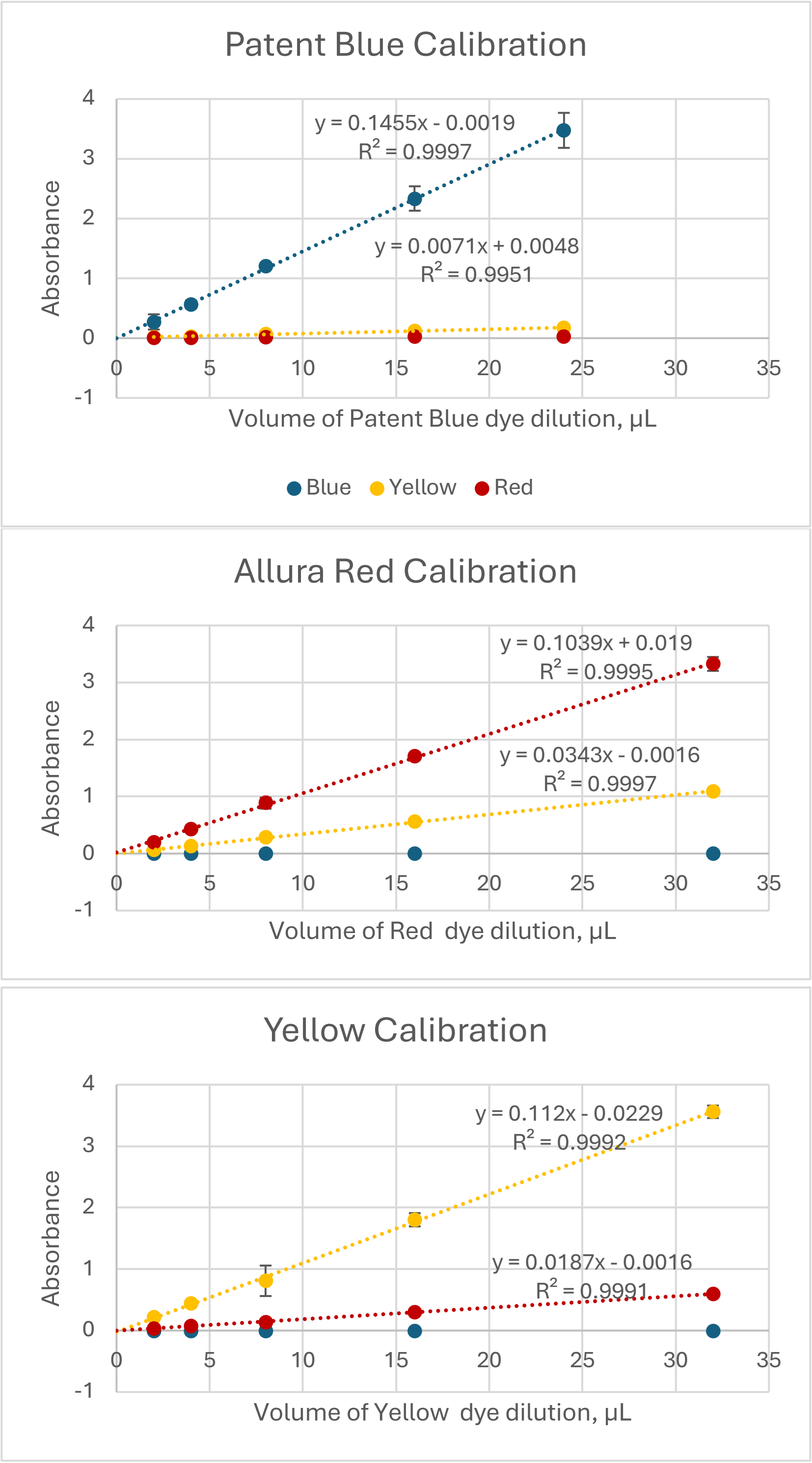

**SI Figure 5 . Electronics schematic.**

The LMNOP-bot is controlled by an Arduino MEGA 2560, with four Pololu A4988 stepper motor drivers. For more information on how to assemble the electronics, please see the LMNOP-bot GitHub.

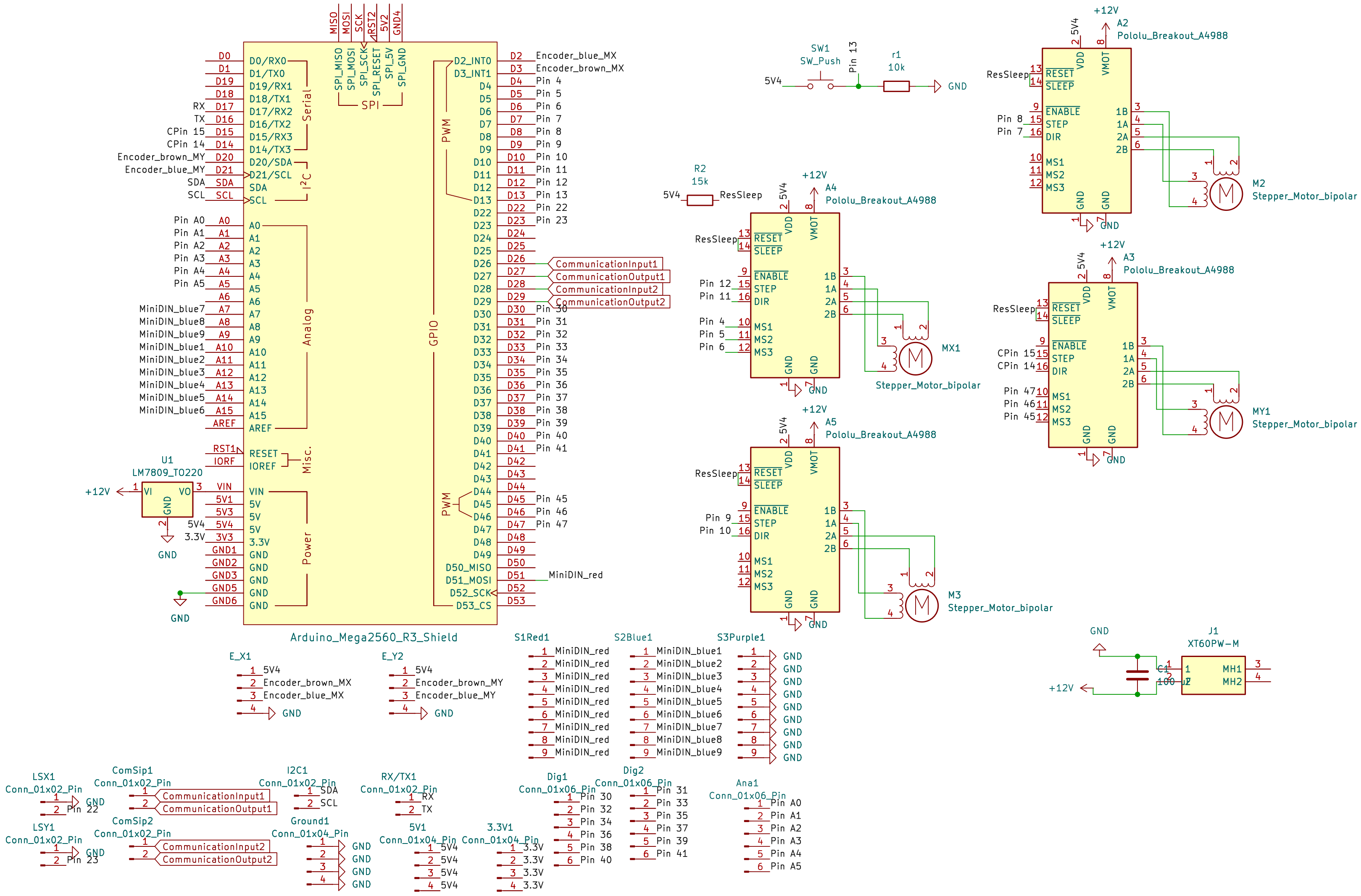

**SI Figure 6**: Sources of vibration, error, and play resolved.

A: inset boxes indicate the location of various sources of vibration and imprecision that were improved.

B: shorter linear bearings had been selected to save money. However, longer linear bearings reduced the play possible between the bearing and the linear shafts and dampened vibrations which had been decreasing repeatability/precision in location.

C: Decreasing the tolerance of the 3D-printed lead screw nuts decreased the imprecision of linear displacement, which could compound over time (see SI Fig 7A).

D: The vibration experienced during normal use could loosen the set screw of the coupler which connects the motor shaft to the lead screw. This could lead to catastrophic uncoupling of the motor from the lead screw, dramatically shifting the puncture location part-way through a run. This was solved by applying LocTite to the threads of the set screw, which both absorbs vibration and increases friction and adhesion of the set screw in place once tightened.

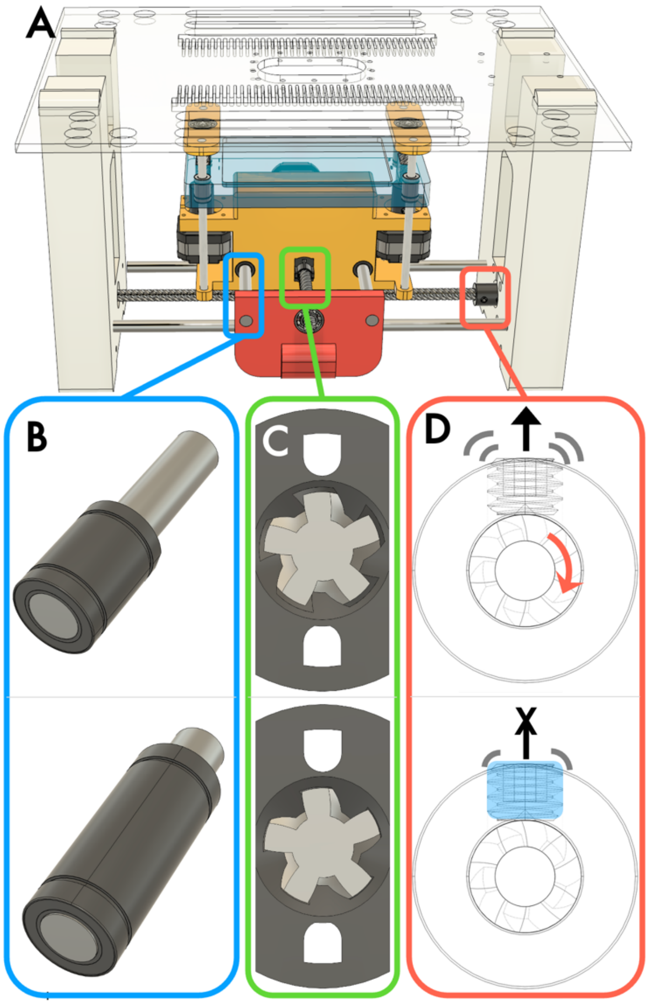


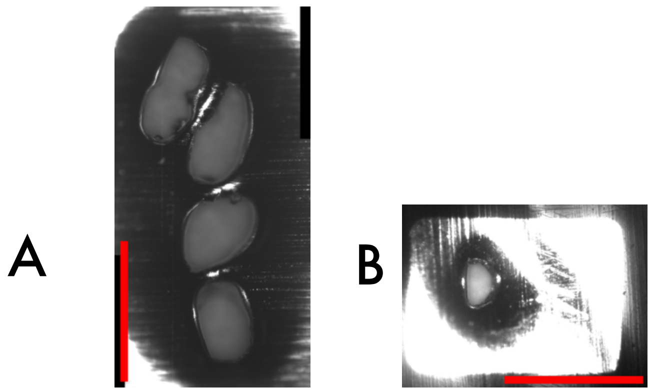


SI Figure 7: example images of 10-hole clusters. Scale bars 1mm. Both collected with zero optional delay.

A: Before addressing sources of vibration and play, we observed significant displacement shifts over the course of 10 hole punctures.

B: After addressing sources of vibration and play, 10 hole clusters often fully overlapped such that the total area was less than the cross sectional area of the needle, as the bevel of the needle leads to single holes having a D-shape, rather than an O-shape.

**SI Figure 8**: Flow response to pressure, for singlet inlet varied from 10 to 25PSI; other 3 inlets held constant at 10 PSI. Use of high-resistance tubing minimizes the impact of varying input pressures, allowing for nearly independent control of flow.

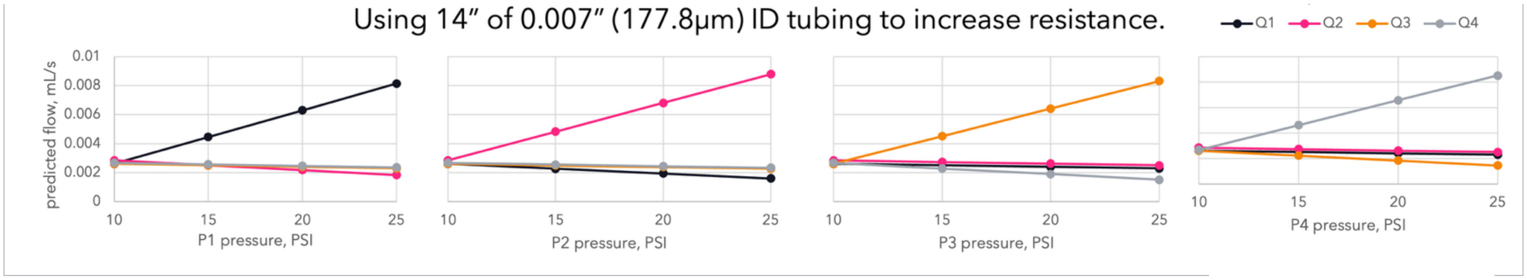

SI Figure 9: Residuals of the fit of measured vs. expected flow proportions ($\phi_m$ vs. $\phi_e$, with trendlines described in Figure 6). A-C, residuals of averaged values ($\bar{y}$); D-F, residuals of individual triplicate points, $y_i$. Run 1 was performed on one day; Run 2 and 3 were performed the next day. It is clear that the yellow flow proportion of run 2 (the first run of the second day) has a different and superior distribution of residuals compared to the other runs. Additionally, the larger and decreasing residuals in runs 1+3 for yellow may have skewed the fit of the Yellow data in such a manner as to impact the Red residuals in the opposite direction, due to their connection by calibration curves.

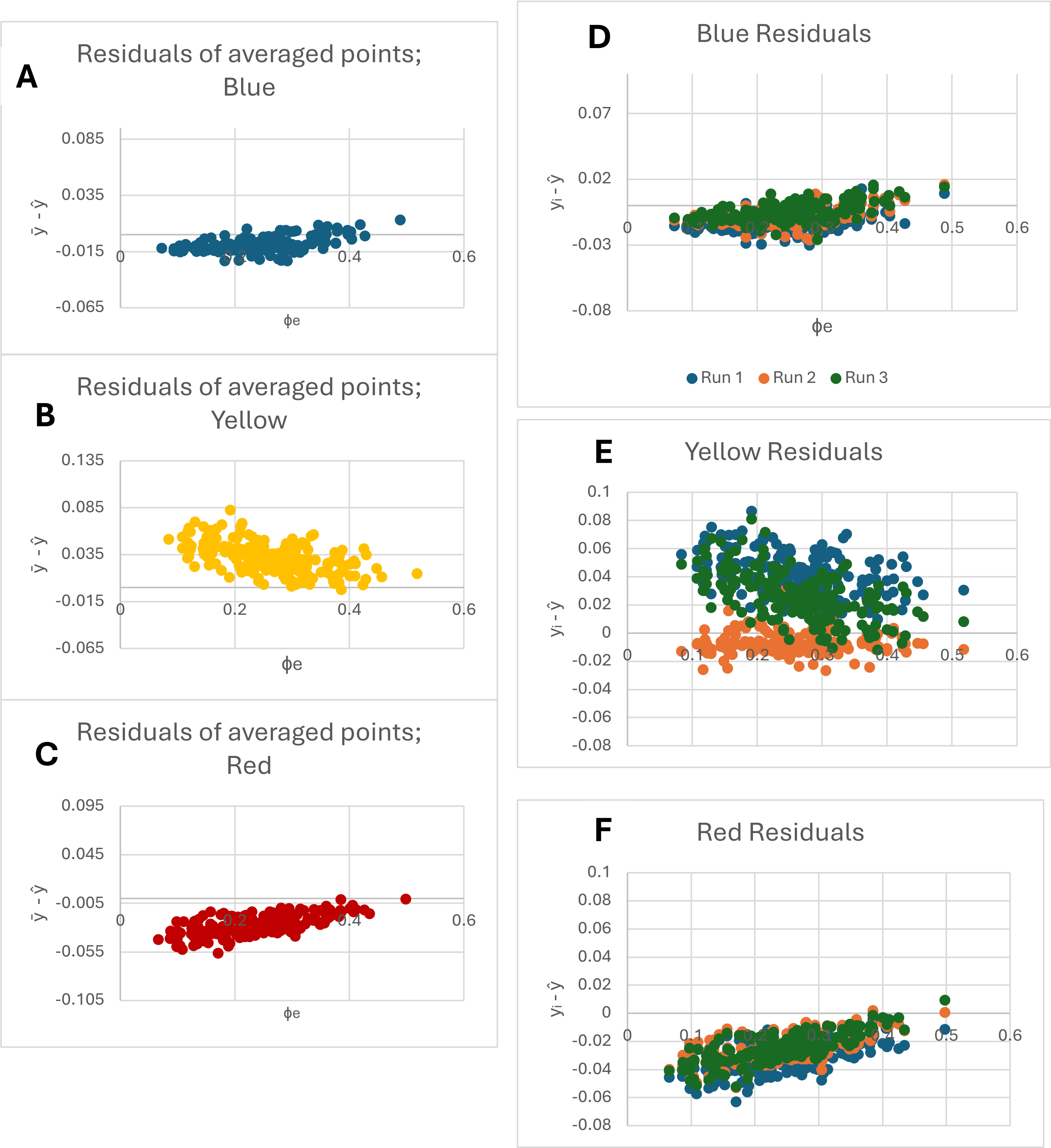
