## Supplemental Arduino Code for "Open-source robotic chip-to-plate interface for high-throughput microfluidic generation of materials libraries"

//This code was written, developed, and documented by Isabel B. Navarro, Gregory Datto, and Lameck Beni.

#include <Wire.h>
#include <SoftwareSerial.h>


//_____________________________________Serial Communication Things________________________________________//
int ArdBaud = 19200;   // Baud rate for the Arduino serial monitor
int AlctBaud = 19200;  // Baud rate on the Alicat (see RS-232 settings), default is 19200

int TxPin = 51;  // Arduino Tx pin, connected to the Alicat Rx RS-232 pin (pin 3 on the 8 pin mini-DIN)
int RxPin = 50;  // Arduino Rx pin, connected to the Alicat Tx RS-232 pin (pin 5 on the 8 pin mini-DIN)

SoftwareSerial regulator_serial(RxPin, TxPin, 1);  // (Arduino Rx pin, Arduino Tx pin, Invert=true).

//____________________________________Pressure/Formulation Variables______________________________________//
const int NUMBER_OF_INPUTS = 4;         // number of inputs (pressure regs / components)
const int NUMBER_OF_FORMULATIONS = 192; // number of formulations.
                                        //max number of formulations on a plate is #wells on the plate / (# collection + 1 purge)
                                        //so if all collections are unique, a 96-well has 48 formulations max
                                        //and a 384-well would have 192 formulations, max

const int NUMBER_OF_COLLECTIONS_PER_FORMULA = 1;    // number of times a given formulation or formulation set is/are collected. 
                                                    //to make the maximum number of  unique formulations, the number is 1.
const int NUMBER_OF_COLLECTIONS_PER_PLATE = NUMBER_OF_COLLECTIONS_PER_FORMULA * NUMBER_OF_FORMULATIONS;

// Pressure array. You can make this list in excel and copy over.
// first dimension is the length of the list, i.e., the number of pressure combinations
// the maximum for this should be # rows x # of columns / (# collections +1)

// the second dimension is the number of inputs/pressure regulators, and therefore, the number of pressures per setting.

// the third dimension is the maximum number of *characters* stored, plus one— that is, the number of digits used PLUS the decimal point, if needed/used, PLUS an unseen return character that marks the end of each entry.
// You do not need to supply the maximum number of digits.
// If you plan to also have an array of times, or increase the plate density/total number of formulations, you may need to decrease the number of digits stored for each setting for the code to fit on the Arduino.
char pressure_array[NUMBER_OF_FORMULATIONS][NUMBER_OF_INPUTS][3] = { 
 
  {"13","11","11","11"}, //the units of pressure are PSI.
  {"17","11","11","11"},
  {"19","11","11","11"},
  {"23","11","11","11"},
  {"11","13","11","11"},
  {"11","17","11","11"},
  {"11","19","11","11"},
  {"11","23","11","11"},
  {"11","11","13","11"},
  {"11","11","17","11"},
  {"11","11","19","11"},
  {"11","11","23","11"},
  {"11","11","11","13"},
  {"11","11","11","17"},
  {"11","11","11","19"},
  {"11","11","11","23"},
  {"11","13","13","13"},
  {"13","13","13","13"},
  {"17","13","13","13"},
  {"19","13","13","13"},
  {"23","13","13","13"},
  {"13","11","13","13"},
  {"13","13","13","13"},
  {"13","17","13","13"},
  {"13","19","13","13"},
  {"13","23","13","13"},
  {"13","13","11","13"},
  {"13","13","13","13"},
  {"13","13","17","13"},
  {"13","13","19","13"},
  {"13","13","23","13"},
  {"13","13","13","11"},
  {"13","13","13","13"},
  {"13","13","13","17"},
  {"13","13","13","19"},
  {"13","13","13","23"},
  {"11","17","17","17"},
  {"13","17","17","17"},
  {"17","17","17","17"},
  {"19","17","17","17"},
  {"23","17","17","17"},
  {"17","11","17","17"},
  {"17","13","17","17"},
  {"17","17","17","17"},
  {"17","19","17","17"},
  {"17","23","17","17"},
  {"17","17","11","17"},
  {"17","17","13","17"},
  {"17","17","17","17"},
  {"17","17","19","17"},
  {"17","17","23","17"},
  {"17","17","17","11"},
  {"17","17","17","13"},
  {"17","17","17","17"},
  {"17","17","17","19"},
  {"17","17","17","23"},
  {"11","19","19","19"},
  {"13","19","19","19"},
  {"17","19","19","19"},
  {"19","19","19","19"},
  {"23","19","19","19"},
  {"19","11","19","19"},
  {"19","13","19","19"},
  {"19","17","19","19"},
  {"19","19","19","19"},
  {"19","23","19","19"},
  {"19","19","11","19"},
  {"19","19","13","19"},
  {"19","19","17","19"},
  {"19","19","19","19"},
  {"19","19","23","19"},
  {"19","19","19","11"},
  {"19","19","19","13"},
  {"19","19","19","17"},
  {"19","19","19","19"},
  {"19","19","19","23"},
  {"11","23","23","23"},
  {"13","23","23","23"},
  {"17","23","23","23"},
  {"19","23","23","23"},
  {"23","23","23","23"},
  {"23","11","23","23"},
  {"23","13","23","23"},
  {"23","17","23","23"},
  {"23","19","23","23"},
  {"23","23","23","23"},
  {"23","23","11","23"},
  {"23","23","13","23"},
  {"23","23","17","23"},
  {"23","23","19","23"},
  {"23","23","23","23"},
  {"23","23","23","11"},
  {"23","23","23","13"},
  {"23","23","23","17"},
  {"23","23","23","19"},
  {"23","23","23","23"},
  {"11","13","17","19"},
  {"13","17","11","19"},
  {"17","11","13","19"},
  {"11","17","13","19"},
  {"13","11","17","19"},
  {"17","13","11","19"},
  {"11","13","19","17"},
  {"13","19","11","17"},
  {"19","11","13","17"},
  {"11","19","13","17"},
  {"13","11","19","17"},
  {"19","13","11","17"},
  {"17","19","11","13"},
  {"19","11","17","13"},
  {"11","17","19","13"},
  {"17","11","19","13"},
  {"19","17","11","13"},
  {"11","19","17","13"},
  {"17","19","13","11"},
  {"19","13","17","11"},
  {"13","17","19","11"},
  {"17","13","19","11"},
  {"19","17","13","11"},
  {"13","19","17","11"},
  {"23","19","17","13"},
  {"19","17","23","13"},
  {"17","23","19","13"},
  {"23","17","19","13"},
  {"19","23","17","13"},
  {"17","19","23","13"},
  {"23","19","13","17"},
  {"19","13","23","17"},
  {"13","23","19","17"},
  {"23","13","19","17"},
  {"19","23","13","17"},
  {"13","19","23","17"},
  {"17","13","23","19"},
  {"13","23","17","19"},
  {"23","17","13","19"},
  {"17","23","13","19"},
  {"13","17","23","19"},
  {"23","13","17","19"},
  {"17","13","19","23"},
  {"13","19","17","23"},
  {"19","17","13","23"},
  {"17","19","13","23"},
  {"13","17","19","23"},
  {"19","13","17","23"},
  {"23","19","17","11"},
  {"19","17","23","11"},
  {"17","23","19","11"},
  {"23","17","19","11"},
  {"19","23","17","11"},
  {"17","19","23","11"},
  {"23","19","11","17"},
  {"19","11","23","17"},
  {"11","23","19","17"},
  {"23","11","19","17"},
  {"19","23","11","17"},
  {"11","19","23","17"},
  {"17","11","23","19"},
  {"11","23","17","19"},
  {"23","17","11","19"},
  {"17","23","11","19"},
  {"11","17","23","19"},
  {"23","11","17","19"},
  {"17","11","19","23"},
  {"11","19","17","23"},
  {"19","17","11","23"},
  {"17","19","11","23"},
  {"11","17","19","23"},
  {"19","11","17","23"},
  {"23","19","13","11"},
  {"19","13","23","11"},
  {"13","23","19","11"},
  {"23","13","19","11"},
  {"19","23","13","11"},
  {"13","19","23","11"},
  {"23","19","11","13"},
  {"19","11","23","13"},
  {"11","23","19","13"},
  {"23","11","19","13"},
  {"19","23","11","13"},
  {"11","19","23","13"},
  {"13","11","23","19"},
  {"11","23","13","19"},
  {"23","13","11","19"},
  {"13","23","11","19"},
  {"11","13","23","19"},
  {"23","11","13","19"},
  {"13","11","19","23"},
  {"11","19","13","23"},
  {"19","13","11","23"},
  {"13","19","11","23"},
  {"11","13","19","23"},
  {"19","11","13","23"}
  };


//_____________________________Variables for taking data from the Regs______________________________________//
char* regulator_addresses[9] = { "AS", "BS", "CS", "DS", "ES", "FS", "GS", "HS", "IS" };

char pressure_settings[NUMBER_OF_INPUTS];
float pressure_settings_float[NUMBER_OF_INPUTS]; // The current pressure values assigned to the pressure regulators

float current_pressures[NUMBER_OF_INPUTS];

int reg_pins[] = { A10, A11, A12, A13, A14, A15, A7, A8, A9 }; //this is our preferred default pin layout; up to 6 pins run from A10-A15, with extras starting at A7

float regulator_max_psis[NUMBER_OF_INPUTS] = { 50, 50, 50, 50 }; // Manually set to the regulators' max psi, please update with correct number and value of settings

float pressure_tolerance = .3; //Units of PSI
float pressure_correction_factor = 1/1.03; //empirically derived correction factor for the relationship between analog voltage out and measured pressure
                                         //If the bot will not move to the next well, typically this is an issue of the pressure not meeting the tolerance. 
                                         //Look at the Serial Monitor (scroll to bottom); look at the pressures measured vs. the pressure settings. The measured pressure(s) are likely are not within the tolerance.
                                         //Next, look at the pressures reported on the regulators' screens (which you labelled). Do they approximately match the readings in your serial monitor?
                                         //If the regulators match the serial monitor and both are outside the tolerance, you may not be supplying enough pressure/flow rate for the regs to quickly adjust and provide pressure. Check your manual valves.
                                         //If the regulator screens are reporting within the tolerance, but the serial monitor does not reflect that, then you should use the difference to estimate a pressure_correction_factor.
                                         //pressure_correction_factor = regulatorScreenPressure / serialPrssureReading


//_____________________________________Z Motor and stuff Things_____________________________________________//

const int DIR_PIN_z1 = 7;
const int STEP_PIN_z1 = 8;
const int DIR_PIN_z2 = 10;
const int STEP_PIN_z2 = 9;

const int z_steppers_up_mem = 450;              
const int z_steppers_up_cas = 450;              
const int z_steppers_needles_just_above = 350;  
const int z_steppers_down = 2;                  
bool move_down_flag = false;

//__________________________________________encoder stuff___________________________________________________//

#define ENCODER_A1 21  // Must be an interrupt-capable pin (2)
#define ENCODER_B1 20  // Must be an interrupt-capable pin (3)

volatile long encoderPositionx = 0;
volatile long global_target_encoderPositionx = 0;
volatile int lastEncodedx = 0;


#define ENCODER_A2 3  // Must be an interrupt-capable pin 2
#define ENCODER_B2 2  // Must be an interrupt-capable pin 3

volatile long encoderPositiony = 0;
volatile long global_target_encoderPositiony = 0;
volatile int lastEncodedy = 0;

//_______________________________________x/y stepper variables_____________________________________________//

int MS1_y = 4;
int MS2_y = 5;
int MS3_y = 6;

int MS1_x = 47;
int MS2_x = 46;
int MS3_x = 45;

const int DIR_PIN_y = 11;
const int STEP_PIN_y = 12;
const int DIR_PIN_x = 14;
const int STEP_PIN_x = 15;

//NOTE: HERE, STEP MEANS LIKE DISTANCE BETWEEN WELL LOCATIONS. 
//there are 4000 encoder counts per rotation, and 1"/rotation. 
//to approximate the encoder counts per step, 
// center-to-center distance (mm) * (0.03937008"/mm) * 4000 encoder counts/inch
//But, it's good to check this empirically before trying to run a chip for the first time.
const int x_encoder_counts_per_step = 709;  //(empirically determined— 1-2 steps higher than calculations predict)
const int y_encoder_counts_per_step = 709;  //for 384-well plate

int x_forward = 1;
int x_back = 0;
int y_forward = 0;
int y_back = 1;
int y_direction;


// REGULAR 96 WELL PLATE
int center_x = 16700;  
int center_y = 17100;  
int center_offset = 0;

const int rising_edge_delay = 30;
int x_delay = 1500;
int y_delay = 1500;
int z_delay = 1500;

const int num_rows = 16;     //8; for 96-well plate
const int num_columns = 24;  //12 for 96-well;
const int num_wells = num_rows * num_columns;

//_____________________________________General workflow variables____________________________________________//

const int buttonPin = 13;
int buttonState = 0;
int moveFlag = 0;
int formulation_counter = 0; // 4/22 set to 0, was -1 (-J)
int collection_counter = 0;
int current_formulation_collection_counter = 0;
int num_collection_events_on_this_plate = 0;
int num_movements = 0;
bool kill_switch = false;
bool equilibrate = true;
int collections_to_be_used = 0;

int up = 1;
int down = 0;

//_________________________________________collect/purge times_______________________________________________//

const long collect_time = 800;  

const long stabilize_time = 500;  


//______________________________________________routines_____________________________________________________//

int previous_location = 0;
int location;

void plateMoveFunc(int end_location) {  //move plate up and down
  if (end_location == 0) {
    location = z_steppers_down;
  } else if (end_location == 1) {
    location = z_steppers_up_mem;
  } else if (end_location == 2) {
    location = z_steppers_up_cas;
  } else {
    location = z_steppers_needles_just_above;
  }
  if (location < previous_location) {
    digitalWrite(DIR_PIN_z1, HIGH);
    digitalWrite(DIR_PIN_z2, HIGH);
    for (int z = 0; z < abs(location - previous_location); z++) {
      PORTH |= (1 << 5) | (1 << 6);
      delayMicroseconds(rising_edge_delay);
      PORTH &= ~((1 << 5) | (1 << 6));
      delayMicroseconds(z_delay);
    }
  } else {
    digitalWrite(DIR_PIN_z1, LOW);
    digitalWrite(DIR_PIN_z2, LOW);
    for (int z = 0; z < abs(location - previous_location); z++) {
      PORTH |= (1 << 5) | (1 << 6);
      delayMicroseconds(rising_edge_delay);
      PORTH &= ~((1 << 5) | (1 << 6));
      delayMicroseconds(z_delay);
    }
  }
  previous_location = location;
}


volatile uint8_t A_x, B_x, lastEncodex, encodedx, sumx;
void updateEncoderX() {
  // Direct pin access for ENCODER_A1 (Pin 21) and ENCODER_B1 (Pin 20)
  // For Pin 21 (PORTF5) and Pin 20 (PORTF4)
  A_x = (PIND & (1 << PD0)) ? HIGH : LOW;  // Read pin D0 (PORTD0) directly
  B_x = (PIND & (1 << PD1)) ? HIGH : LOW;  // Read pin D0 (PORTD1) directly

  encodedx = (A_x << 1) | B_x;
  sumx = (lastEncodedx << 2) | encodedx;

  if (sumx == 0b0001 || sumx == 0b0111 || sumx == 0b1110 || sumx == 0b1000) {
    encoderPositionx++;
  } else if (sumx == 0b0010 || sumx == 0b1101 || sumx == 0b1011 || sumx == 0b0100) {
    encoderPositionx--;
  }
  lastEncodedx = encodedx;
}

volatile uint8_t A_y, B_y, lastEncodey, encodedy, sumy;

void updateEncoderY() {
  // Direct pin access for ENCODER_A2 (Pin 2) and ENCODER_B2 (Pin 3)
  // For Pin 2 (PORTD2) and Pin 3 (PORTD3)
  A_y = (PINE & (1 << PINE4)) ? HIGH : LOW;  // Read pin 2 (PORTE4) directly
  B_y = (PINE & (1 << PINE5)) ? HIGH : LOW;  // Read pin 3 (PORTE5) directly

  encodedy = (A_y << 1) | B_y;
  sumy = (lastEncodedy << 2) | encodedy;

  if (sumy == 0b0001 || sumy == 0b0111 || sumy == 0b1110 || sumy == 0b1000) {
    encoderPositiony++;
  } else if (sumy == 0b0010 || sumy == 0b1101 || sumy == 0b1011 || sumy == 0b0100) {
    encoderPositiony--;
  }
  lastEncodedy = encodedy;
}

int current_microstep_setting_x = 0;
float steps_per_rotation_x = 200;
float encoder_counts_per_step_x = 20;

void changeStepX(int microstep_size) {
  if (microstep_size == 0) {  //full step
    digitalWrite(MS1_x, LOW);
    digitalWrite(MS2_x, LOW);
    digitalWrite(MS3_x, LOW);
    x_delay = 1500;
    steps_per_rotation_x = 200;
    encoder_counts_per_step_x = 20;
    current_microstep_setting_x = 1;
  } else if (microstep_size == 1) {  //half step
    digitalWrite(MS1_x, HIGH);
    digitalWrite(MS2_x, LOW);
    digitalWrite(MS3_x, LOW);
    x_delay = 1500;
    steps_per_rotation_x = 400;
    encoder_counts_per_step_x = 10;
    current_microstep_setting_x = 2;
  } else if (microstep_size == 2) {  //quarter step
    digitalWrite(MS1_x, LOW);
    digitalWrite(MS2_x, HIGH);
    digitalWrite(MS3_x, LOW);
    x_delay = 1500;
    steps_per_rotation_x = 800;
    encoder_counts_per_step_x = 5;
    current_microstep_setting_x = 3;
  }
}

void moveStepperX(long number_of_encoder_counts, int dir) {
  changeStepX(0);
  delay(10);

  unsigned long previous_encoder_value;

  if (dir == 0) {
    global_target_encoderPositionx = global_target_encoderPositionx - number_of_encoder_counts;
    while (abs(global_target_encoderPositionx - encoderPositionx) > 13) {
      int fudge_steps = (int)((((float)(encoderPositionx - global_target_encoderPositionx)) / 4000.0) * steps_per_rotation_x);
      if (fudge_steps < 0) {
        digitalWrite(DIR_PIN_x, LOW);
        for (int x = 0; x < abs(fudge_steps); x++) {
          digitalWrite(STEP_PIN_x, HIGH);
          delayMicroseconds(rising_edge_delay);
          digitalWrite(STEP_PIN_x, LOW);
          delayMicroseconds(x_delay);
          if (abs(global_target_encoderPositionx - encoderPositionx) <= 13) {
            break;
          }
        }
      } else {
        digitalWrite(DIR_PIN_x, HIGH);
        for (int x = 0; x < abs(fudge_steps); x++) {
          digitalWrite(STEP_PIN_x, HIGH);
          delayMicroseconds(rising_edge_delay);
          digitalWrite(STEP_PIN_x, LOW);
          delayMicroseconds(x_delay);
          if (abs(global_target_encoderPositionx - encoderPositionx) <= 13) {
            break;
          }
        }
      }
      if (abs(global_target_encoderPositionx - encoderPositionx) <= 13) {
        break;
      }
      while ((abs(global_target_encoderPositionx - encoderPositionx) < (encoder_counts_per_step_x * 5)) && (current_microstep_setting_x < 3)) {
        changeStepX(current_microstep_setting_x);
      }
    }
  } else {
    global_target_encoderPositionx = global_target_encoderPositionx + number_of_encoder_counts;
    while (abs(global_target_encoderPositionx - encoderPositionx) > 13) {
      int fudge_steps = (int)((((float)(encoderPositionx - global_target_encoderPositionx)) / 4000.0) * steps_per_rotation_x);
    
      if (fudge_steps < 0) {
        digitalWrite(DIR_PIN_x, LOW);
        for (int x = 0; x < abs(fudge_steps); x++) {
          digitalWrite(STEP_PIN_x, HIGH);
          delayMicroseconds(rising_edge_delay);
          digitalWrite(STEP_PIN_x, LOW);
          delayMicroseconds(x_delay);
          if (abs(global_target_encoderPositionx - encoderPositionx) <= 13) {
            break;
          }
        }
      } else {
        digitalWrite(DIR_PIN_x, HIGH);
        for (int x = 0; x < abs(fudge_steps); x++) {
          digitalWrite(STEP_PIN_x, HIGH);
          delayMicroseconds(rising_edge_delay);
          digitalWrite(STEP_PIN_x, LOW);
          delayMicroseconds(x_delay);
          if (abs(global_target_encoderPositionx - encoderPositionx) <= 13) {
            break;
          }
        }
      }
      if (abs(global_target_encoderPositionx - encoderPositionx) <= 13) {
        break;
      }
      while ((abs(global_target_encoderPositionx - encoderPositionx) < (encoder_counts_per_step_x * 5)) && (current_microstep_setting_x < 3)) {
        changeStepX(current_microstep_setting_x);
      }
    }
  }
}

int current_microstep_setting_y = 0;
float steps_per_rotation_y = 200;
float encoder_counts_per_step_y = 20;

void changeStepY(int microstep_size) {
  if (microstep_size == 0) {
    digitalWrite(MS1_y, LOW);
    digitalWrite(MS2_y, LOW);
    digitalWrite(MS3_y, LOW);
    y_delay = 1000;
    steps_per_rotation_y = 200;
    encoder_counts_per_step_y = 20;
    current_microstep_setting_y = 1;
  } else if (microstep_size == 1) {
    digitalWrite(MS1_y, HIGH);
    digitalWrite(MS2_y, LOW);
    digitalWrite(MS3_y, LOW);
    y_delay = 1000;
    steps_per_rotation_y = 400;
    encoder_counts_per_step_y = 10;
    current_microstep_setting_y = 2;
  } else if (microstep_size == 2) {
    digitalWrite(MS1_y, LOW);
    digitalWrite(MS2_y, HIGH);
    digitalWrite(MS3_y, LOW);
    y_delay = 1000;
    steps_per_rotation_y = 800;
    encoder_counts_per_step_y = 5;
    current_microstep_setting_y = 3;
  }
}

void moveStepperY(int number_of_encoder_counts, int dir) {
  changeStepY(0);
  unsigned long previous_encoder_value;
  delay(10);
  if (dir == 0) {
    global_target_encoderPositiony = global_target_encoderPositiony - number_of_encoder_counts;
    while (abs(global_target_encoderPositiony - encoderPositiony) > 13) {
      int fudge_steps = (int)((((float)(encoderPositiony - global_target_encoderPositiony)) / 4000.0) * steps_per_rotation_y);
      if (fudge_steps < 0) {
        digitalWrite(DIR_PIN_y, LOW);
        for (int y = 0; y < abs(fudge_steps); y++) {
          digitalWrite(STEP_PIN_y, HIGH);
          delayMicroseconds(rising_edge_delay);
          digitalWrite(STEP_PIN_y, LOW);
          delayMicroseconds(y_delay);
          if (abs(global_target_encoderPositiony - encoderPositiony) <= 13) {
            break;
          }
        }
      } else {
        digitalWrite(DIR_PIN_y, HIGH);
        for (int y = 0; y < abs(fudge_steps); y++) {
          digitalWrite(STEP_PIN_y, HIGH);
          delayMicroseconds(rising_edge_delay);
          digitalWrite(STEP_PIN_y, LOW);
          delayMicroseconds(y_delay);
          if (abs(global_target_encoderPositiony - encoderPositiony) <= 13) {
            break;
          }
        }
      }
      if (abs(global_target_encoderPositiony - encoderPositiony) <= 13) {
        break;
      }
      while ((abs(global_target_encoderPositiony - encoderPositiony) < (encoder_counts_per_step_y * 5)) && (current_microstep_setting_y < 3)) {
        changeStepY(current_microstep_setting_y);
      }
    }
  } else {
    global_target_encoderPositiony = global_target_encoderPositiony + number_of_encoder_counts;
    while (abs(global_target_encoderPositiony - encoderPositiony) > 13) {
      int fudge_steps = (int)((((float)(encoderPositiony - global_target_encoderPositiony)) / 4000.0) * steps_per_rotation_y);
      if (fudge_steps < 0) {
        digitalWrite(DIR_PIN_y, LOW);
        for (int y = 0; y < abs(fudge_steps); y++) {
          digitalWrite(STEP_PIN_y, HIGH);
          delayMicroseconds(rising_edge_delay);
          digitalWrite(STEP_PIN_y, LOW);
          delayMicroseconds(y_delay);
          if (abs(global_target_encoderPositiony - encoderPositiony) <= 13) {
            break;
          }
        }
      } else {
        digitalWrite(DIR_PIN_y, HIGH);
        for (int y = 0; y < abs(fudge_steps); y++) {
          digitalWrite(STEP_PIN_y, HIGH);
          delayMicroseconds(rising_edge_delay);
          digitalWrite(STEP_PIN_y, LOW);
          delayMicroseconds(y_delay);
          if (abs(global_target_encoderPositiony - encoderPositiony) <= 13) {
            break;
          }
        }
      }
      if (abs(global_target_encoderPositiony - encoderPositiony) <= 13) {
        break;
      }
      while ((abs(global_target_encoderPositiony - encoderPositiony) < (encoder_counts_per_step_y * 5)) && (current_microstep_setting_y < 3)) {
        changeStepY(current_microstep_setting_y);
      }
    }
  }
}

void formatPressureSettingMessage(char* buffer, const char* prefix, const char* value) {  //make pressures in a format that can be read
  memset(buffer, 0, sizeof(buffer));
  strcpy(buffer, prefix);  // Copy the prefix
  strcat(buffer, value);   // Concatenate the value to the buffer
  strcat(buffer, "\r");    // Append the carriage return
}

void waitForEquilibrium(float* pressure_settings_float) {  // took out A2 and 3
  Serial.println(F("Hello! Waiting for equilibrium"));
  unsigned long entered_time = millis(); //Start the clock for stabilize time

  for (int i = 0; i < NUMBER_OF_INPUTS; i++) { //Read in the pressure from analog pins and use to set current_pressures
    current_pressures[i] = analogRead(reg_pins[i]) * (5.0 / 1023.0) * (regulator_max_psis[i] / 5.0) * pressure_correction_factor;
  }

  for (int i = 0; i < NUMBER_OF_INPUTS; i++) { //Print the results for troubleshooting
    Serial.print(current_pressures[i], 3);
    Serial.print(F(","));
  }

  Serial.println();

  //if you're not at equilibrium and haven't pushed the kill switch, keep reading in the pressures
  while (!atEquilibrium(pressure_settings_float, current_pressures) && (!kill_switch)) { 
    for (int i = 0; i < NUMBER_OF_INPUTS; i++) {
      current_pressures[i] = analogRead(reg_pins[i]) * (5.0 / 1023.0) * (regulator_max_psis[i] / 5.0) * pressure_correction_factor;
    }
    
    for (int i = 0; i < NUMBER_OF_INPUTS; i++) { //print out current pressures
      Serial.print(current_pressures[i], 3);
      Serial.print(F(","));
    }
    Serial.print(F(","));

    for (int i = 0; i < NUMBER_OF_INPUTS; i++) { //and the intended set pressures, for troubleshooting
      Serial.print(pressure_settings_float[i], 3);
      Serial.print(F(","));
    }
    Serial.println();

    if (digitalRead(buttonPin) == HIGH) { //if you happen to push the kill switch here, set pressures to zero
      kill_switch = true;      
      setZeroPressure();
    }
  }
}

bool atEquilibrium(float* pressure_settings_float, float* current_pressures) {
  for (int i = 0; i < NUMBER_OF_INPUTS; i++) {
    if (abs(pressure_settings_float[i] - current_pressures[i]) > pressure_tolerance) {
      return false;
    }    
  }
  return true;
}

/**
 * Sends pressures to regulators 
 */
void setPressures(char pressures[][3], bool use_delay = true) {  
  
  for (int i = 0; i < NUMBER_OF_INPUTS; i++) {
    char buffer[50];
    formatPressureSettingMessage(buffer, regulator_addresses[i], pressures[i]);  
    regulator_serial.print(buffer);
    pressure_settings_float[i] = atof(pressures[i]);
    pressure_settings[i] = pressures[i];
  }

  if (use_delay) {
    delay(50);
  }
  
  waitForEquilibrium(pressure_settings_float);

  for (int i = 0; i < NUMBER_OF_INPUTS; i++) {
    current_pressures[i] = atof(pressures[i]);    
  }

}

void setZeroPressure() {
  char zero_pressures[NUMBER_OF_INPUTS][3]; 
  
  for (int i = 0; i < NUMBER_OF_INPUTS; i++) {
    strcpy(zero_pressures[i], "0");
  }

  setPressures(zero_pressures); 
}

void waitForTime(unsigned long duration) {
  unsigned long start = millis();
  while (((millis() - start) < duration) && !kill_switch) {
    if (digitalRead(buttonPin) == HIGH) {
      kill_switch = true;
      setZeroPressure();
    }
  }
}

void equilibrize() {  //function that makes the pressures set in waste wells    
  setPressures(pressure_array[formulation_counter], false); 
  waitForTime(stabilize_time);
}

void center() {
  moveStepperX(center_x - center_offset, 0);  
  moveStepperY(center_y - center_offset, 0);  
}

void homeSteppers() {
  if (global_target_encoderPositiony > (-100)) {
    moveStepperY(global_target_encoderPositiony + 100, 0);  
  } else {
    moveStepperY(abs(global_target_encoderPositiony) - 100, 1);  
  }
  if (global_target_encoderPositionx > (-100)) {
    moveStepperX(global_target_encoderPositionx + 100, 0);  
  } else {
    moveStepperX(abs(global_target_encoderPositionx) - 100, 1);  
  }
}

void moveToNewLocation() {

  // Checking if *not* at end of row (numbered), in which case, move in x (along column)
  if (((num_movements + 1) % num_columns) != 0) {
    moveStepperY(y_encoder_counts_per_step, y_direction);
  }

  // If at end of row, are we at the end of the plate?
  else if ((num_movements + 1) == num_wells) {
    moveStepperX(x_encoder_counts_per_step, x_back);
  }

  // If at end of row and not end of plate, reverse x direction and move in y (along row)
  else if (((num_movements + 1) / num_columns) % 2 == 0) {
    y_direction = y_back;
    moveStepperX(x_encoder_counts_per_step, x_forward);
  } else {
    y_direction = y_forward;
    moveStepperX(x_encoder_counts_per_step, x_forward);
  }
  //Serial.println();
  Serial.println(num_movements);
  num_movements++;
}

void setup() {
  Serial.begin(ArdBaud);              // Arduino Serial monitor interface
  regulator_serial.begin(AlctBaud);   // Alicat Serial interface (make sure baud rate matches Alicat setting)
  
  while (!Serial); // if serial connection is not made, do not proceed

  pinMode(ENCODER_A1, INPUT_PULLUP);
  pinMode(ENCODER_B1, INPUT_PULLUP);
  attachInterrupt(digitalPinToInterrupt(ENCODER_A1), updateEncoderX, CHANGE);
  attachInterrupt(digitalPinToInterrupt(ENCODER_B1), updateEncoderX, CHANGE);
  pinMode(ENCODER_A2, INPUT_PULLUP);
  pinMode(ENCODER_B2, INPUT_PULLUP);
  attachInterrupt(digitalPinToInterrupt(ENCODER_A2), updateEncoderY, CHANGE);
  attachInterrupt(digitalPinToInterrupt(ENCODER_B2), updateEncoderY, CHANGE);

  setZeroPressure();

  pinMode(buttonPin, INPUT);  //turn on all pins
  pinMode(DIR_PIN_x, OUTPUT);
  pinMode(STEP_PIN_x, OUTPUT);
  pinMode(DIR_PIN_y, OUTPUT);
  pinMode(STEP_PIN_y, OUTPUT);
  pinMode(STEP_PIN_z1, OUTPUT);
  pinMode(DIR_PIN_z1, OUTPUT);
  pinMode(STEP_PIN_z2, OUTPUT);
  pinMode(DIR_PIN_z2, OUTPUT);
  pinMode(MS1_x, OUTPUT);
  pinMode(MS2_x, OUTPUT);
  pinMode(MS3_x, OUTPUT);
  changeStepX(0);
  pinMode(MS1_y, OUTPUT);
  pinMode(MS2_y, OUTPUT);
  pinMode(MS3_y, OUTPUT);
  changeStepY(0);
  digitalWrite(STEP_PIN_z1, HIGH);
  digitalWrite(STEP_PIN_z2, HIGH);

  while (digitalRead(buttonPin) == LOW);  //wait for button press

  encoderPositionx = 0;
  encoderPositiony = 0;
}

void loop() {
  // Serial.println(encoderPositionx); // this can be used to empirically check/estimate first well location and well spacings.
  // Serial.println(encoderPositiony);
  // Serial.println("------------");
  while (digitalRead(buttonPin) == LOW);  // wait for user to start
  
  center();                               // centers over first well
  formulation_counter = 0;
  
  while (digitalRead(buttonPin) == LOW);  // wait for user to continue

  plateMoveFunc(1);                       // move steppers to membrane (z dir)
  
  while (digitalRead(buttonPin) == HIGH); // ensure button is low before continuing: avoid inadvertly triggering killswitch
  
  y_direction = y_back;  
  
  equilibrize();        // come to your first set of pressures
  plateMoveFunc(3);     // retract plate away from needle

  // while we haven't finished collecting...
  while (collection_counter < (NUMBER_OF_COLLECTIONS_PER_PLATE)) {    
    current_formulation_collection_counter = 0;
    num_movements = 0;

    while (true) {
      // make all collections for the current formulation
      while (true) {
        moveToNewLocation();
        plateMoveFunc(2);  //bring needles into casette
        waitForTime(collect_time);
        plateMoveFunc(3);                          //bring needles to just above the plate
        collection_counter++;                      //increase the total number of collection events by 1
        current_formulation_collection_counter++;  //increase the total number of collection events for a single formulation by 1
        
        if (collection_counter == NUMBER_OF_COLLECTIONS_PER_PLATE) {  //run out of wells to collect into
          break;
        }
        if (current_formulation_collection_counter == NUMBER_OF_COLLECTIONS_PER_FORMULA) {  //run out of collections for a given set of particles
          current_formulation_collection_counter = 0;
          break;
        }
        if (kill_switch) {
          break;
        }
      }

      moveToNewLocation();
      plateMoveFunc(1);  //bring needles to membrane level

      if (kill_switch) {  //break after move so that you are in a clean well
        break;
      }
      if (collection_counter == NUMBER_OF_COLLECTIONS_PER_PLATE) {  //break the loop if you are using the last well
        break;
      }
      if (formulation_counter == (NUMBER_OF_FORMULATIONS - 1)) {  // when you run out of pressure settings for a given input change inputs
        formulation_counter = 0;
        // no break?
      } else {
        formulation_counter++;
        equilibrize();
      }
      plateMoveFunc(3);  //bring needles to just above the plate
    }

    if (kill_switch) {
      break;
    } 

    if (collection_counter == NUMBER_OF_COLLECTIONS_PER_PLATE) {
      break;
    }
  }

  formulation_counter = 0;
  setZeroPressure();
  plateMoveFunc(0);
  homeSteppers();
  kill_switch = false;
  while (true);
}
